# Active surface waves drive rippling in *Myxococcus xanthus* colonies

**DOI:** 10.1101/2025.11.28.691037

**Authors:** Aaron R. Bourque, Peter A. E. Hampshire, Ricard Alert, Joshua W. Shaevitz

## Abstract

During periods of predation or starvation, populations of the gliding bacterium *Myxococcus xanthus* self-organize into striking wave-like structures termed ripples. This phenomenon was thought to arise from wave collisions triggering synchronized reversals of cell motility. However, using three-dimensional microscopy, we find no evidence for such synchronization during rippling. Instead, we show that ripples are surface waves with a period of ∼ 20 min, wavelength of ∼ 100 *µ*m and an amplitude of 6 to 20 cell widths at the top of a thick film of cells, akin to surface waves seen in fluids. We propose a physical model of rippling as surface waves of an active nematic liquid crystal. Two key predictions of this model are verified experimentally: the rippling wavelength increases with the surface tension at the film–air interface, and it decreases with substrate stiffness, which regulates the availability of water coating the bacterial film. These findings reveal the physical basis of rippling and highlight the role of active surface waves in shaping collective biological behavior.

## I. INTRODUCTION

Cell populations exhibit a wide range of morphologies and collective behaviors, such as aggregation and directed motion, often in response to environmental cues. These processes span multiple length scales and are typically driven by complex biochemical or mechanical interactions between individual agents. For instance, certain bacterial species form structured surface-attached communities, or biofilms, by adhering to substrates and secreting polymeric substances that recruit other cells [1]. The slime mold *Dictyostelium discoideum* aggregates through chemotaxis, guided by self-generated gradients of diffusible chemoattractants, ultimately forming multicellular slugs and fruiting bodies [2, 3]. In contrast, the soil bacterium *Myxococcus xanthus* forms multicellular streams and layered aggregates through short-range mechanical interactions, without relying on long-range chemical signaling [4–9].

*M. xanthus* is a surface-gliding bacterium that exhibits a range of social behaviors throughout its life cycle, making it a model organism for studying multicellular structure formation in response to environmental cues [10]. Colonies of these rod-shaped cells spread as a thin film across a substrate when nutrients are abundant, and aggregate into mound-shaped fruiting bodies under starvation conditions [11, 12]. Recent work has shown that aggregation is driven by changes in single-cell motility, specifically through modulation of speed and reversal frequency [7]. Mound formation is initiated when the population organizes into an active nematic liquid crystal, where topological defects promote the emergence of three-dimensional, droplet-like fruiting bodies [8, 13].

During both major phases of its life cycle, *M. xanthus* can form wave-like patterns in a process known as rippling that has fascinated scientists for decades [14–18]. During predation, a colony can form ripples as it swarms over prey such as bacteria or yeast [19, 20]. Similarly, during starvation, ripples frequently emerge in the regions between developing fruiting bodies [14, 17, 21].

The prevailing model for rippling in *M. xanthus* proposes that coordination of cellular reversals, mediated by contact-dependent signaling, drives the formation of periodic density fluctuations known as accordion waves [17]. In this model, cells reverse direction synchronously at high local densities, particularly at wave crests, creating alternating regions of high and low cell density that propagate and reflect from one another [15]. However, key aspects of this model remain untested. Evidence for synchronized reversals and their coincidence with wave crest collisions comes from experiments at very low cell densities, where colonies form a monolayer with gaps [17]. This contrasts with the multilayered populations typically used in rippling studies [5, 7, 22, 23], where cell density is practically uniform. Moreover, the accordion-wave model predicts that the rippling wavelength is set by the distance that a cell travels before undergoing a collision-induced reversal, and is therefore determined directly by motility parameters. While previous work found the observed wavelength to be quantitatively consistent with the accordion-wave model, this was only tested for one wavelength and condition, without varying motility [18].

Here, we measure ripples in multilayered *M. xanthus* colonies on different substrates using 3D imaging and cell tracking, and we show that the accordion-wave model is inconsistent with several of our observations. We find that ripples are standing surface waves at the top of a thick, dense population of cells. Cells move and reverse throughout the population, but we do not observe synchronization of reversals in space or time. Motivated by these findings, we propose a physical model in which rippling arises as surface waves on an active nematic film. We test this model by varying the surface tension of the bacterial film using a surfactant, and by employing hydrogel substrates of different compositions and concentrations. The changes in wavelength produced by these perturbations cannot be explained by variations in cell speed or reversal frequency, but are qualitatively captured by the active surface wave model.

## II. RESULTS

### A. Ripples are standing surface waves

We first sought to characterize the three-dimensional (3D) morphology of rippling colonies. When *M. xanthus* is subjected to nutrient-poor conditions, rippling waves emerge throughout the colony once it reaches a thickness of several cell layers and begins expanding into the surrounding environment. In circular colonies, we observe circumferential wavefronts, consistent with previous observations [14] (Fig. 1a, Supp. Video 1).

To investigate the 3D structure of ripples, we used a laser-scanning confocal profilometer (Keyence VK-X1000; see Methods and Materials) that measures reflectance at different heights to localize the top surface of a sample. We observe that ripples are height undulations on the surface of a thick colony of cells (Fig. 1b). Wave amplitudes range from 3 to 10 *µ*m, corresponding to approximately six to twenty cell layers. Ripples form on top of a basal film of cells with a height of approximately 4 to 10 *µ*m (Fig. 1b, see Supplementary Information Sections I and II). We recorded a time series of height maps of rippling colonies to examine their dynamics (Supp. Video 2). Clear height oscillations are observed both at fixed spatial positions (Fig. 1c) and in spatial profiles taken along the wave vector direction (orthogonal to the orientation of the wave crests; Fig. 1d). In these profiles, we identify locations with minimal amplitude variation (nodes) and others with maximal variation (antinodes; Fig. 1d). Taken together, these observations show that ripples are standing surface waves.

**FIG. 1.**
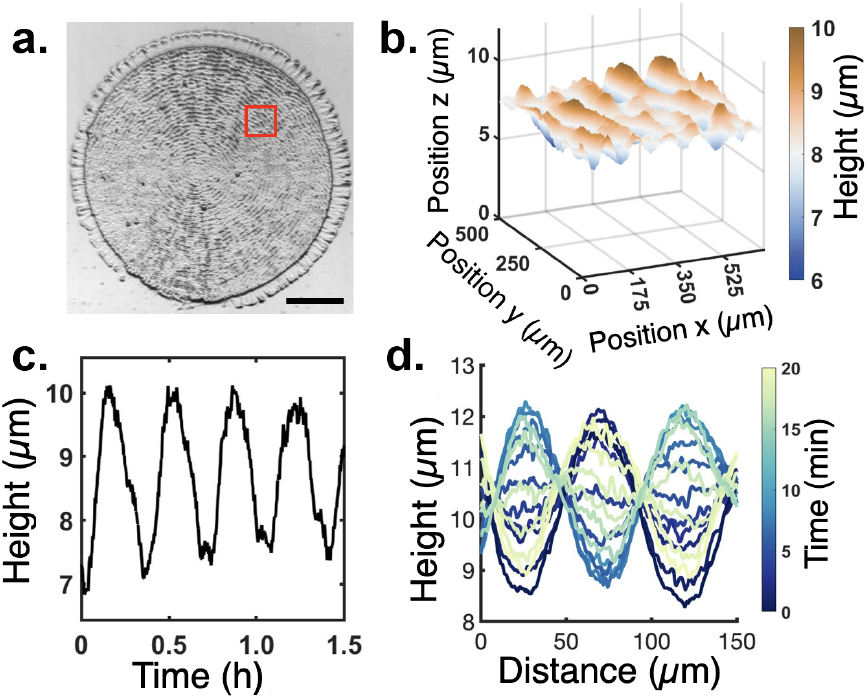
Rippling in *Myxococcus xanthus* produces standing surface waves. (a) Brightfield image of rippling on the surface of a circular *M. xanthus* colony showing circumferential wavefronts. Scale bar = 2.5 mm. (b) Topographical height map of the boxed region in panel (a). A height of zero corresponds to the underlying substrate, indicating that ripples are undulations on the top surface of a thick film of cells. (c) Height of a single point in the colony as a function of time, showing periodic oscillations. (d) Temporal progression of the height profile along a radial line within the colony shown in (a), demonstrating standing wave behavior with static nodes and antinodes.

### B. Cell reversals are not synchronized

To investigate individual cell behavior within ripples, we sparsely inoculated *M. xanthus* cells expressing either green or red fluorescent proteins into a culture of unlabeled cells (0.5% ratio; see Methods and Materials), and imaged them using spinning disc confocal microscopy (Fig. 2a). This approach allows us to track the dynamics of single cells within a large, dense population. We focused our imaging near the minima of the rippling undulations, where single cells remain within the focal plane over time. We tracked cells over several wave periods (Fig. 2b; see Materials and Methods). Cells move primarily parallel to the local wave vector, along the radial direction and orthogonal to the circular wave fronts, and frequently reverse direction by 180°. From these trajectories, we extracted the average reversal period (7.0 ± 0.4 min), average cell speed (4.3 *µ*m/min), and the positions and times of individual reversals.

**FIG. 2.**
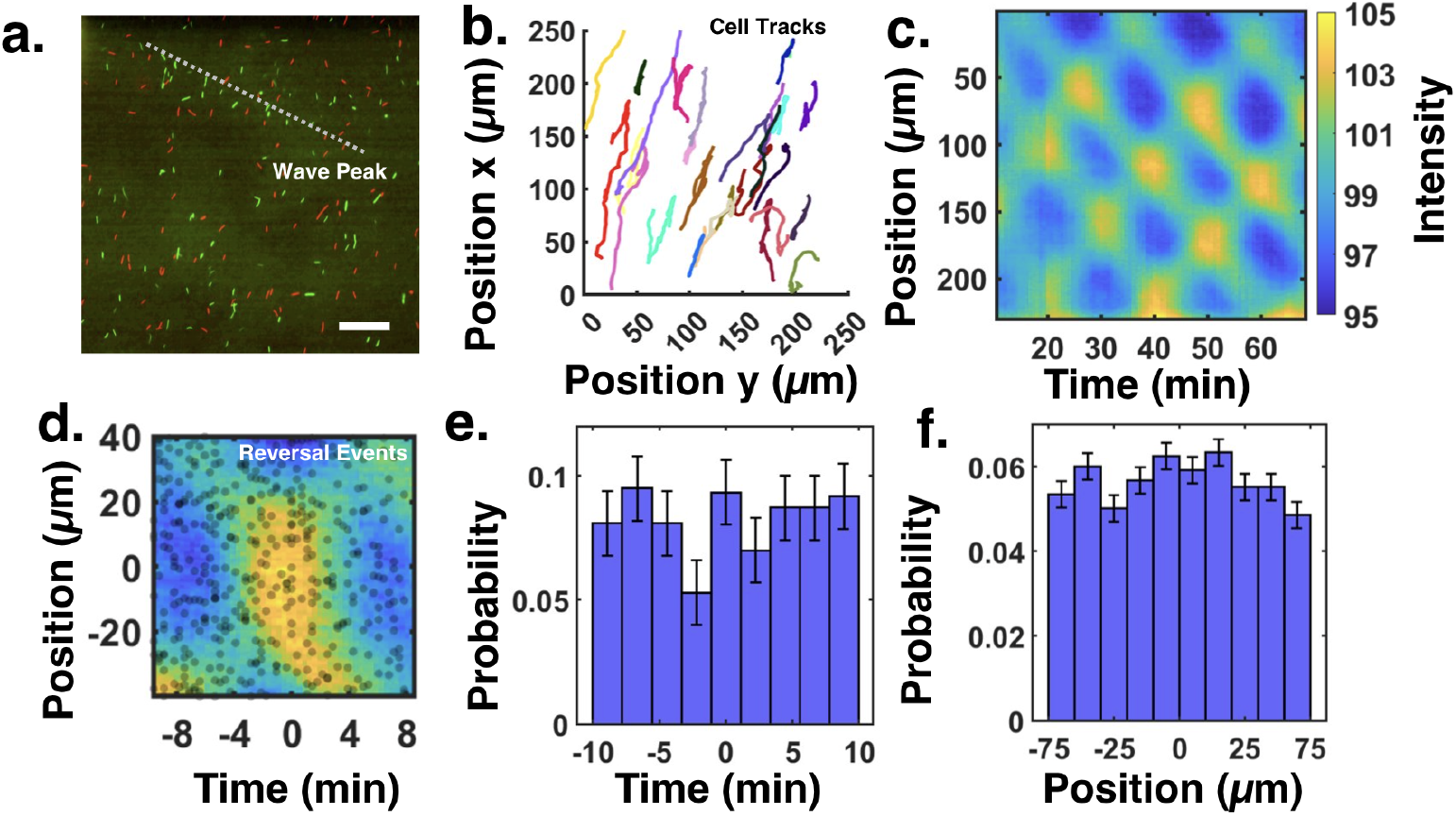
Cell reversals are not correlated with wave peaks. (a) Image of sparsely labeled fluorescent cells within ripples. Images are acquired at the bottom plane of the rippling surface. Both red and green fluorescent labels were used to increase the density of trackable single cells. The dashed gray line indicates the wave peak analyzed in panel (c). (b) A subset of cell tracks from the movie corresponding to panel (a). Each track, individually colored, is a trajectory from a labeled cell with a duration of 45 minutes. Cells move roughly along the wave vector and undergo frequent 180° directional reversals. (c) Kymograph of the background autofluorescence intensity of unlabeled cells. This background autofluorescence serves as a proxy for projected cell density, which reveals the surface waves. The kymograph is obtained by averaging the autofluorescence intensity along the dashed line in (a) at each time point. (d) Autofluorescence intensity averaged around each occurrence of a wave peak, yellow spots in panel (b). Black dots indicate the time and position of cell reversal events relative to their nearest wave peak. (e,f) Probability distribution of elapsed time (e) and distance (f) between the occurrence of a wave peak and the nearest cell reversals. Error bars indicate one standard deviation Poisson uncertainties, computed as the square root of the number of counts in each bin. A Kolmogorov-Smirnov test fails to reject a null hypothesis that the distribution in (f) corresponds to a uniform distribution (*p* < 0.001).

To test the accordion-wave model, we investigated the correlation between cell reversal events and the surface waves. The unlabeled cells in each image, which greatly outnumber the labeled cells, produce a level of green autofluorescence that serves as a readout of the local colony thickness. A kymograph of this fluorescence intensity shows the characteristic pattern of a standing wave, with nodes and antinodes and constant wavelength and period (Fig. 2c; see Methods). The peaks in the kymograph correspond to ripple crests, previously described as wave collisions but now identified as the antinodes of a standing wave.

A key prediction of the accordion-wave model is that cell reversal events are synchronized in space and time and coincide with wave peaks. We averaged the autofluorescence signal within a fixed spatiotemporal region centered on each wave peak across all experiments (Fig. 2d). We then plotted the positions and times of all cell reversal events relative to their nearest wave peak in space and time (Fig. 2d, black points). The distribution of reversal events is nearly uniform around wave peaks in both time (Fig. 2e) and space (Fig. 2f), indicating that cell reversals are uncorrelated with wave peaks. We do not observe global synchronization of cell reversals, which would appear as a sharp clustering of events around wave peaks in both space and time [18]. A Kolmogorov–Smirnov test of the spatial distribution in Fig. 2f supports uniformity (*p* < 0.001). Likewise, the time intervals between reversal events and wave peaks show no peak around the onset of peak formation (Fig. 2e). These observations are inconsistent with the accordion-wave model.

### C. Theory of rippling as active surface waves

Motivated by the lack of agreement with the accordion-wave model, we developed a new model of rippling in *M. xanthus* films. We propose that ripples are surface waves of an active nematic liquid crystal [24–26]. Previous work showed that single layers of *M. xanthus* form an extensile active nematic [8, 13]. We now extend this model to consider the dynamics of the upper surface of a thick multilayered colony.

To capture the physical mechanism of the waves, we propose a minimal model of the cell colony as an incompressible active nematic, of which we consider a two-dimensional semi-infinite vertical cross-section (Fig. 3a).

**FIG. 3.**
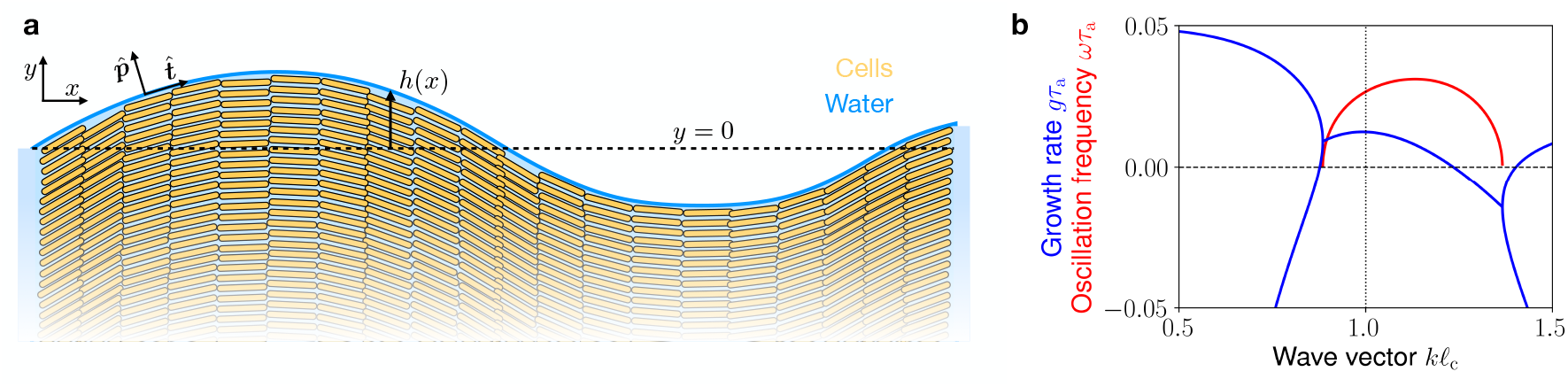
Surface waves on an active nematic. (a) Schematic of the model. *M. xanthus* cells (yellow) produce active stresses that deform the water-air interface at the colony’s surface (blue). (b) Dispersion relation of the active surface waves, showing both the growth rate and the oscillation frequency of perturbations about the flat-interface, homogeneous state (*θ* = 0, ***v*** = **0**, *h* = 0). Rates are scaled by the active time *τ*_a_ = *η*/*ζ*, and the wave vector is scaled with the capillary length 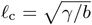 (see text). Parameter values are specified in Table I of the SI.

Force balance imposes

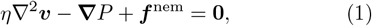

where *η* is the colony’s viscosity, ***v*** is the velocity field, and *P* is the pressure that enforces the incompressibility condition **∇** *·* ***v*** = 0. The nematic force density is given by ***f*** ^nem^ = **∇** *·* ***σ***^nem^ and arises from distortions in the nematic director 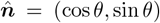, which defines the local axis of cell alignment. The nematic stress ***σ***^nem^ includes an active stress ***σ***^a^ = −*ζ****Q***, with coefficient *ζ* > 0 and proportional to the nematic tensor *Q*_*αβ*_ = *n*_*α*_*n*_*β*_ − *δ*_*αβ*_/2. It also includes flow-alignment and antisymmetric stresses (Supplementary Information Section IX.A). In addition, the total stress includes viscous stress and pressure, 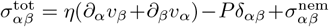, in terms of which force balance is simply **∇** *·* ***σ***^tot^ = 0.

At the colony-air interface, at height *h*(*x, t*), we impose two boundary conditions: (i) no shear stress: 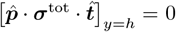, and (ii) normal stress balance with surface tension *γ* and a restoring force *bh* due to the water meniscus that surrounds the bacterial film [9]: 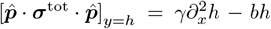. Here, we approximated the curvature of the interface as 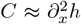, and the unit vectors 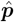 and 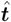 refer to the directions perpendicular and tangential to the surface, respectively (Fig. 3a). The gravity-like restoring force *bh* captures the energy penalty of extracting water from the underlying hydrogel substrate (Supplementary Information Section IX.C). Moreover, the film’s surface moves with the flow, which can be expressed as ∂_*t*_*h* = *v*_*y*_(*y* = *h*) − *v*_*x*_(*y* = *h*)∂_*x*_*h*.

From the physics of liquid crystals [27], the dynamics of the director field 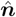 follow

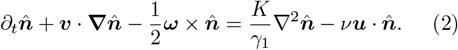

The left-hand side describes how the director is advected by the flow and co-rotated by its vorticity ***ω*** = **∇** × ***v***. The right-hand side accounts for nematic elasticity, which tends to relax the director distortions, where *K* is the Frank elastic constant and *γ*_1_ is the rotational viscosity, and for flow alignment (with coefficient *ν*) under shear described by the symmetric part of the strain-rate tensor *u*_*αβ*_ = (∂_*α*_*v*_*β*_ + ∂_*β*_*v*_*α*_)/2.

Following previous work [25, 26], we study the emergence of waves by performing a linear stability analysis of the flat-interface, homogeneous state (*θ* = 0, ***v*** = **0**, *h* = 0). The linearized dynamics of the height *h* and director angle at the interface *θ*(*y* = 0) are given by (Supplementary Information Section IX.D)

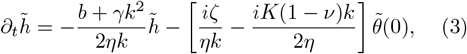

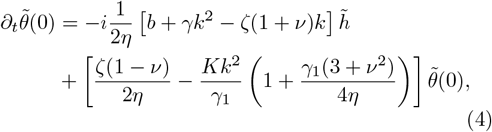

where tildes indicate the components with wavenumber *k* of a Fourier transform along the *x* axis and we have used the shorthand that *f* (*y*) := *f* (*k, y, t*) for any function *f*. These dynamics encode the fact that a director distortion produces active flows, which drive interfacial fluctuations. In turn, these interfacial fluctuations are resisted by surface tension and the restoring force, and the resulting flows rotate the director. This feedback gives rise to surface waves [26]. Below, we discuss how they compare with our experimental observations.

The analysis above predicts that the growth rate of oscillating interfacial perturbations is *g* = [*ζ*(1 − *ν*) − (*b* + *γk*^2^)/*k*)] /(4*η*) when neglecting the contribution of nematic elasticity (*K* = 0, see Supplementary Information Section IX.E, Fig. 3b). Hence, the flat interface is unstable if *ζ >* 2*γ/* [𝓁_c_(1 − *ν*)], where 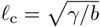 is the capillary length of the interface. Thus, the emergence of waves depends on the competition of active and capillary stresses, encoded in the active capillary number [28, 29] Ca_A_ ≡ *ζ*𝓁_c_*/γ*. The most unstable mode occurs for *k* = 1/𝓁_c_. Hence, we predict that the waves have a selected wavelength *λ*_s_ = 2*π*𝓁_c_. The corresponding frequency can be expressed as 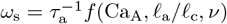, where *τ*_a_ = *η/ζ* and 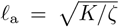 are the active time and length of the liquid crystal, and *f* is a function of three dimensionless parameters (Supplementary Information Section IX.E). Overall, surface waves emerge with a wavelength controlled by passive interfacial properties, encoded in the capillary length 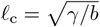, and a period that scales with the active time *τ*_a_ = *η*/*ζ*.

Finally, we use these results to estimate parameter values (see Supplementary Note Section IX.F). Given that the observed wavelengths are of the order *λ* ∼ 100 *µ*m (Fig. 1d), we estimate the capillary length as 𝓁_c_ ≈ *λ/*(2*π*) ∼ 15 *µ*m. With a measured surface tension of the cell medium *γ* ≈ 65 mN/m (Fig. 4a–b), and taking *ν* ≈ − 1.1 so that the colony has flow-aligning behavior [27], we estimate *ζ* ∼ 4 kPa for waves to emerge. Next, by requiring that there are oscillations at the selected wavelength, we estimate *K* ∼ 2 kPa *µ*m^2^, for which we took *γ*_1_ = *η* for simplicity. Finally, comparing the predicted oscillation frequency to the measured wave period *T* ∼ 20 min (Fig. 1c,d), we estimate *η* ∼ 300 Pa min for the viscosity of the bacterial film, which gives an active time *τ*_a_ ∼ 5 s.

**FIG. 4.**
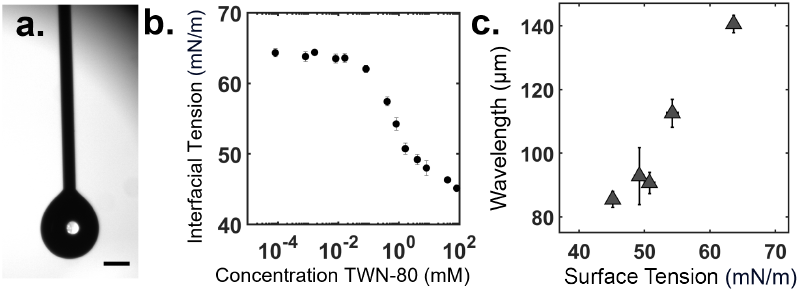
The rippling wavelength is modulated by the addition of surfactant. (a) Example image of the pendant-drop technique to measure surface tension. A droplet of medium is suspended from a 1 mm needle attached to a reservoir. For water, the measured surface tension is 73 ± 1 mN/m. (b) Surface tension of growth medium (CTTYE) containing different concentrations of the surfactant Tween-80. The surface tension of CTTYE without any added surfactant is measured to be 63.6 ± 0.3 mN/m. (c) Measured rippling wave-length of *M. xanthus* colonies as a function of surface tension.

### D. The rippling wavelength increases with surface tension

Our model predicts that the rippling wavelength is controlled by capillary forces at the film’s surface. To test this prediction, we added the surfactant Tween-80 to the substrate and the cell medium and repeated the rippling experiments. We first used the pendant drop method to measure the surface tension of the medium for increasing surfactant concentrations (Fig. 4a) [30]. In this technique, a drop of medium is held at the end of a syringe, and it is imaged via brightfield microscopy. We use the OpenDrop software to fit the droplet shape and calculate the shape factor, a dimensionless parameter that describes how gravity and surface tension balance in a pendant or sessile drop [31]. This factor is obtained by numerically solving the Young–Laplace equation following the approach of Berry et al. [30]. We validated our measurements using water as a control and find that the addition of surfactant decreases the surface tension from approximately 65 to 45 mN/m (Fig. 4b). Consistent with our surface-wave model, we find that the rippling wavelength increases with the surface tension of the medium (Fig. 4c). This dependence cannot be explained by the accordion-wave model, which does not involve surface tension.

### E. The rippling wavelength decreases with substrate stiffness

Our model also predicts that the rippling wavelength depends on the properties of the underlying hydrogel substrate, encoded in the interfacial restoring force *bh*. Recent work showed that, on porous water-filled substrates, bacterial cells are surrounded by a meniscus of water extracted from the substrate [9]. These menisci produce capillary forces that attract cells to each other and push cells onto the substrate. Extracting water from the substrate has an energetic cost [9]; hence, substrate properties influence capillary forces and, in turn, should affect rippling.

To test the role of the substrate on rippling, we varied substrate composition, changing both the gelling agent and its concentration (Fig. 5a). We used agarose with a concentration varying between 0.5% and 1.5%, which is a typical range used for experiments, and phytagel, which is an optically clear hydrogel alternative to agarose. We found that higher concentration of gelling agent, which yields stiffer substrates, results in smaller rippling wavelengths (Fig. 5a). This trend is consistent with our model: A substrate with a higher concentration of polymer can bind more water, which implies a higher energetic cost of extracting water from it, and hence a higher coefficient *b* of the restoring force, which yields a smaller wavelength. To further validate this picture, we measured the decay length of the meniscus around individual cells [9]. For the different substrates that we used, the rippling wavelength correlates with the meniscus decay length (Fig. 5b).

**FIG. 5.**
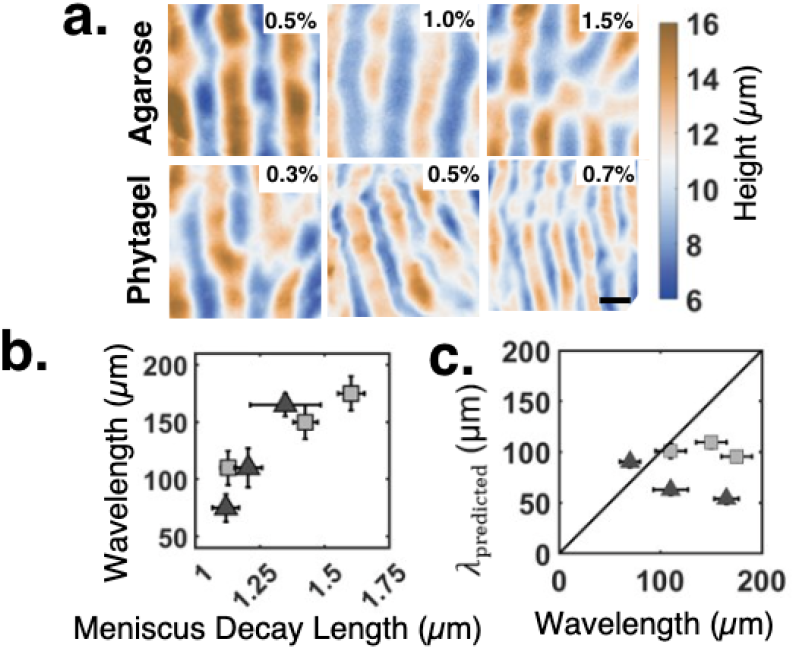
The rippling wavelength is modulated by substrate properties. (a) Height fields of rippling on hydrogels made from three different concentrations of agarose (top) and phytagel (bottom). (b) The measured rippling wavelength increases with the meniscus decay length (see Methods). Data from both agarose (triangles) and phytagel (squares) is shown. Error bars represent standard error of the mean. (c) The wavelength predicted by the accordion-wave model, *λ*_predicted_ = 2(*vτ* + *δ*), does not agree with the measured rippling wavelength. Data from agarose (triangles) and phytagel (squares) substrates is shown. A line of slope 1, which would indicate model agreement, is shown for comparison.

In addition to water extraction, changing substrate properties can influence both individual and collective motility in *M. xanthus* [32–35]. Single cells move at different speeds on agar of varying concentration [35], and colonies of *M. xanthus* expand faster on stiffer substrates [32, 34]. In the accordion-wave model, the wavelength is given by the distance traveled by cells before collision-induced reversals,

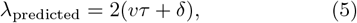

where *v* is the cell speed, *τ* is the average time between reversals, and *δ* is the width of the wave crest. Thus, sub-strate properties could in principle also affect the rippling wavelength within the framework of the accordion-wave model by modulating cell motility itself.

To test this possibility, we performed single-cell tracking on each of the substrates to measure cell motility parameters and compare the predicted wavelength with the observed rippling wavelengths. Our results indicate that the substrate has a mild effect on motility, and that the predicted wavelength does not match the observed rippling wavelength (Fig. 5c). Taken together, our findings indicate that changing substrate stiffness affects rippling through capillary phenomena, as qualitatively captured by our model.

## DISCUSSION

Our results recast rippling in *Myxococcus xanthus* as an interfacial phenomenon: surface waves driven by active stresses on the colony surface. Three findings motivate this view. First, three dimensional profiling reveals standing surface undulations with static nodes and antinodes that overlay a thick, dense film. Second, single-cell tracking shows frequent reversals throughout the colony without coordination in space or time, which is inconsistent with models that rely on synchronized, collision-induced reversals. Third, the wavelength is controlled by physical parameters of the film’s interface. The wavelength increases with surface tension and decreases with substrate polymer concentration, which reduces water availability. These trends are captured by an active nematic model in which active stresses due to cell motility drive surface waves against the restoring effects of surface tension and the resistance of water to be extracted from the hydrogel substrate. Interestingly, our model predicts that the rippling wavelength is set by the interface capillary length, which is a passive property, whereas active stresses set the oscillation period. Future work could address the roles of the film’s thickness, substrate friction, the permeation of water through the hydrogel substrate, and nonlinearities in controlling wave properties.

The new view of rippling as an interfacial phenomenon has several implications. It reveals that physical principles of active matter and capillarity underlie the long-studied bacterial collective behavior of rippling. It also suggests practical ways to manipulate the colony’s morphology by tuning the medium’s surface tension, humidity, osmolarity, or substrate properties. More broadly, whereas active surface waves were theoretically predicted and experimentally found in other systems [24–26, 36– 41], such as colloidal chiral fluids [37] and cytoskeletal filament-motor suspensions [25, 40], our results show their relevance for a naturally-occurring biological process. Similar active surface waves could shape the interfaces of other biological systems such as biomolecular condensates [42], bacteria-containing droplets [41], dense microbial films [43], and epithelial tissues [38, 44] — all of which exhibit a competition between internal active stresses and interfacial forces. Understanding these processes in a quantitative, physical framework offers a route to predict and control how active materials sculpt their own boundaries, with implications ranging from biofilm architecture to the design of self-organizing soft interfaces.

## METHODS AND MATERIALS

### A. Sample Preparation

*M. xanthus* DK16122 cells are grown in CTTYE (1% casitone, 10mM Tris–HCl, 1 mM KH2PO4, and 8mM MgSO4 at pH 7.6) overnight to a concentration corresponding to a range of OD600 ≈ 0.4–0.8, where OD600 is the optical density at a wavelength of 600nm. This culture is then concentrated and resuspended into CF (1 mM KH2PO4, 8 mM MgSO4, 0.02% (NH4)2SO4, 0.2% citrate, 0.2% pyruvate, and 150 mg/liter peptone at pH of 7.6) to an OD600 value of 2. Then, 10*µ*l of the concentrated culture is placed onto a CF plate of agarose or phytagel substrate, and it is allowed to dry uncovered for a period of 3 hours. The sample is then submerged in 5 mL of CTTYE medium and incubated at 32C for a period of 4 hours. The liquid media is then removed, and the samples are dried by flame for an hour, after which colonies begin to ripple. For surface tension modulation experiments, TWN-80 dissolved in medium is added at various concentrations to the substrate and submersion medium.

### B. Imaging

Height measurements are recorded using a Keyence VK-X1000 microscope. Images of the colony surface are acquired by scanning a laser onto the sample, and recording the reflectance of the laser light. After scanning the sample at various heights, the height field of the sample is determined from the maximum reflectance for each pixel. Imaging was done at a frame rate between 1.0 and 4.0 min^−^1 for 1 to 8 hours. A 20x magnification air objective is used (NA = 0.36), and the field of view of the rippling region is ∼500×650 µm.

Single cells were imaged using spinning disc confocal microscopy with a Nikon Ti2 inverted microscope with Yokogawa W1 and SoRa Module (W1). *M. xanthus* strains DK10547 and LS3908 were cultured and inoculated in the wild-type culture above in a 1:200 ratio. Images of the fluorescent cells were acquired at a frame rate of 2.0 min^−^1 for 45 minutes to 2 hours. The field of view of the rippling region is ∼260×260 µm.

### C. Data Analysis

#### 1. Wavelength Measurement

The 2D Fourier Transform was calculated on mean-centered height map data collected from the Keyence VK-X1000 microscope by calculating the fast Fourier Transform for the rows and columns of the image of size M × N. This results in the power spectrum of the image, an M × N-sized matrix in which each element represents spatial frequencies in *k*-space. The power spectrum is shifted such that the DC component of the signal is at the center of the matrix, resulting in two peaks that represent the dominant spatial frequency composing the waves (see Supplementary Information Section V). To calculate the rippling wavelength from this, we calculate the static structure factor by averaging the signal of the power spectrum of identical values of *k*. A log-normal distribution is then fitted for each time point of the duration of the rippling behavior. The mean of the fitted curve is determined to be the rippling wavelength for the image.

#### 2. Cell Tracking

To track cells within waves, shot noise is first removed from fluorescent confocal images using the Denoise software from Nikon NIS-Elements. Individual cells are then segmented using an adaptive threshold based on local mean intensity around each pixel of the image and then using the regionprops MATLAB function. Tracks are constructed by solving the linear assignment problem of all of the located centroids of the image as in Jaqaman et al. [45]. Reversal instances are detected by calculating the autocorrelation of instantaneous velocities of the cell tracks. When the autocorrelation value of the velocity changes in sign, a reversal is recorded at the midpoint time between the two frames.

## ACKNOWLEDGMENTS

We thank the Confocal Imaging Facility, a Nikon Center of Excellence, in the Department of Molecular Biology at Princeton University for instrument use and technical advice. We thank Matthew Black, Katherine Copenhagen, Endao Han, Ahmed Al Harraq, Allie Zhao, Nicholas O’Reilly, and Cassidy Yang for their assistance in the research and many useful discussions. R.A. thanks Aparna Baskaran, Paarth Gulati, Frank Jülicher, John D. McEnany, Ned S. Wingreen, and Qiwei Yu for discussions. P.H. and R.A. thank Katherine Copenhagen for providing the estimate of the defect density in *M. xanthus* monolayers. This work was supported by the NSF through awards PHY-1806501 and PHY2210346, and the Center for the Physics of Biological Function (PHY-1734030). R.A. acknowledges funding from the European Union through ERC Starting Grant “Living Fluctuations” (No. 101114584).

## AUTHOR CONTRIBUTIONS

A.B. performed the experiments and analyzed the data. J.W.S. supervised the study. R.A. and P.H. developed the model. P.H. performed the theory calculations. All authors interpreted the results and wrote the manuscript.

## References

[1] L. Hall-Stoodley, J. W. Costerton, and P. Stoodley, Bacterial biofilms: from the Natural environment to infectious diseases, Nature Reviews Microbiology 2, 95 (2004).

[2] Y. Murata and T. Ohnishi, Dictyostelium discoideum fruiting bodies observed by scanning electron microscopy, Journal of Bacteriology 141, 956 (1980).

[3] F. Siegert and C. J. Weijer, Spiral and concentric waves organize multicellular Dictyostelium mounds, Current Biology 5, 937 (1995).

[4] O. Sozinova, Y. Jiang, D. Kaiser, and M. Alber, A three-dimensional model of myxobacterial aggregation by contact-mediated interactions, Proceedings of the National Academy of Sciences 102, 11308 (2005).

[5] Y. Zhang, A. Ducret, J. W. Shaevitz, and T. Mignot, From individual cell motility to collective behaviors: insights from a prokaryote, Myxococcus xanthus, FEMS Microbiology Reviews 36, 149 (2012).

[6] R. Balagam and O. A. Igoshin, Mechanism for collective cell alignment in Myxococcus xanthus bacteria, PLOS Computational Biology 11, e1004474 (2015).

[7] G. Liu, A. Patch, F. Bahar, D. Yllanes, R. D. Welch, M. C. Marchetti, S. Thutupalli, and J. W. Shaevitz, Self-driven phase transitions drive Myxococcus xanthus fruiting body formation, Physical Review Letters 122, 248102 (2019).

[8] K. Copenhagen, R. Alert, N. S. Wingreen, and J. W. Shaevitz, Topological defects promote layer formation in Myxococcus xanthus colonies, Nature Physics 17, 211 (2021).

[9] M. E. Black, C. Fei, R. Alert, N. S. Wingreen, and J. W. Shaevitz, Capillary interactions drive the self-organization of bacterial colonies, Nature Physics 21, 1444 (2025).

[10] R. Mohr, W. McLaughlin, Y. Xiao, S. Chen, K. Kowallis, S. Walsh, J. Liu, and et al., Diversity of Myxococcus xanthus rippling behaviors is controlled by the frz chemotaxis pathway, mBio 9, e01754 (2018).

[11] M. Dworkin, Nutritional regulation of morphogenesis in Myxococcus xanthus, Journal of Bacteriology 86, 67 (1963).

[12] J. H. Parish, K. R. Wedgwood, and D. G. Herries, Morphogenesis in Myxococcus xanthus and Myxococcus virescens (myxobacterales), Archives of Microbiology 107, 343 (1976).

[13] E. Han, C. Fei, R. Alert, K. Copenhagen, M. D. Koch, N. S. Wingreen, and J. W. Shaevitz, Local polar order controls mechanical stress and triggers layer formation in Myxococcus xanthus colonies, Nature Communications 16, 952 (2025).

[14] L. J. Shimkets and D. Kaiser, Induction of coordinated movement of Myxococcus xanthus cells, Journal of Bacteriology 152, 451 (1982).

[15] O. A. Igoshin, A. Goldbeter, D. Kaiser, and G. Oster, A biochemical oscillator explains several aspects of Myxococcus rippling, Proceedings of the National Academy of Sciences 98, 14913 (2001).

[16] A. Stevens and L. Søgaard-Andersen, Making waves: pattern formation by a cell-surface-associated signal, Trends in Microbiology 13, 249 (2005).

[17] O. Sliusarenko, J. Neu, D. R. Zusman, and G. Oster, Accordion waves in Myxococcus xanthus, Proceedings of the National Academy of Sciences 103, 1534 (2006).

[18] H. Zhang, Z. Vaksman, D. B. Litwin, P. Shi, H. B. Kaplan, and O. A. Igoshin, The mechanistic basis of Myxococcus xanthus rippling behavior and its physiological role during predation, PLoS Computational Biology 8, e1002715 (2012).

[19] J. E. Berleman, J. Scott, T. Chumley, and J. R. Kirby, Predataxis behavior in Myxococcus xanthus, Proceedings of the National Academy of Sciences 105, 17127 (2008).

[20] J. E. Berleman, T. Chumley, P. Cheung, and J. R. Kirby, Rippling is a predatory behavior in myxococcus xanthus, Journal of Bacteriology 188, 5888 (2006).

[21] R. D. Welch and D. Kaiser, Cell behavior in traveling wave patterns of myxobacteria, Proceedings of the National Academy of Sciences 98, 14907 (2001).

[22] S. Thiery and C. Kaimer, The predation strategy of Myxococcus xanthus, Frontiers in Microbiology 10.3389/fmicb.2020.00002 (2020).

[23] J. Pèrez, A. Moraleda-Muñoz, F. J. Marcos-Torres, and J. Muñoz-Dorado, Bacterial predation: 75 years and counting!, Environmental Microbiology 18, 766 (2016).

[24] H. Soni, W. Luo, R. A. Pelcovits, and T. R. Powers, Stability of the interface of an isotropic active fluid, Soft Matter 15, 6318 (2019).

[25] R. Adkins, I. Kolvin, Z. You, S. Witthaus, M. C. Marchetti, and Z. Dogic, Dynamics of active liquid interfaces, Science 377, 768 (2022).

[26] P. Gulati, F. Caballero, I. Kolvin, Z. You, and M. C. Marchetti, Traveling waves at the surface of active liquid crystals, Soft Matter 20, 7703 (2024).

[27] P. G. de Gennes and J. Prost, The Physics of Liquid Crystals (Oxford University Press, 1993).

[28] L. Giomi, Spontaneous division and motility in active nematic droplets, Physical Review Letters 112, 147802 (2014).

[29] R. Alert, Fingering instability of active nematic droplets, Journal of Physics A: Mathematical and Theoretical 55, 234009 (2022).

[30] J. D. Berry, M. J. Neeson, R. R. Dagastine, D. Y. C. Chan, and R. F. Tabor, Measurement of surface and interfacial tension using pendant drop tensiometry, Journal of Colloid and Interface Science 454, 226 (2015).

[31] J. D. Berry, Opendrop: Open-source droplet tracking and analysis software, https://github.com/jdber1/opendrop (2024), accessed: 2025-11-03.

[32] N. Faiza, R. Welch, and A. Patteson, Substrate stiffness modulates collective colony expansion of the social bacterium Myxococcus xanthus, APL Bioengineering 9, 016104 (2025).

[33] M. E. Asp, M.-T. Ho Thanh, D. A. Germann, R. J. Carroll, A. Franceski, R. D. Welch, A. Gopinath, and A. E. Patteson, Spreading rates of bacterial colonies depend on substrate stiffness and permeability, PNAS Nexus 1, pgac025 (2022).

[34] L. J. Ritchie, E. R. Curtis, K. A. Murphy, and R. D. Welch, Profiling Myxococcus xanthus swarming phenotypes through mutation and environmental variation, Journal of Bacteriology 203, e00306.

[35] W. Shi and D. R. Zusman, The two motility systems of Myxococcus xanthus show different selective advantages on various surfaces, Proceedings of the National Academy of Sciences of the United States of America 90, 3378 (1993).

[36] S. Sankararaman and S. Ramaswamy, Instabilities and waves in thin films of living fluids, Physical Review Letters 102, 118107 (2009).

[37] V. Soni, E. S. Bililign, S. Magkiriadou, S. Sacanna, D. Bartolo, M. J. Shelley, and W. T. M. Irvine, The odd free surface flows of a colloidal chiral fluid, Nature Physics 15, 1188 (2019).

[38] C.-L. Lv, Z.-Y. Li, S.-D. Wang, and B. Li, Morphodynamics of interface between dissimilar cell aggregations, Communications Physics 7, 351 (2024).

[39] L. Langford and A. K. Omar, Phase separation, capillarity, and odd-surface flows in chiral active matter, Physical Review Letters 134, 068301 (2025).

[40] A. Sciortino, H. A. Faizi, D. A. Fedosov, L. Frechette, P. M. Vlahovska, G. Gompper, and A. R. Bausch, Active membrane deformations of a minimal synthetic cell, Nature Physics 21, 799 (2025).

[41] K. Chang, Y. Li, M. Yuan, M. Sano, Z. You, and H. P. Zhang, Collective bacterial motion drives interfacial waves and shape dynamics in phase-separated droplets, (2025).

[42] F. Raßhofer, S. Bauer, A. Ziepke, I. Maryshev, and E. Frey, Capillary wave formation in conserved active emulsions, 26 (2025).

[43] S. Liu, Y. Li, Y. Wang, and Y. Wu, Emergence of largescale mechanical spiral waves in bacterial living matter, Nature Physics 20, 1015 (2024).

[44] C. L. Lv and B. Li, Interface morphodynamics in living tissues, Soft Matter 21, 3670 (2025).

[45] K. Jaqaman, D. Loerke, M. Mettlen, H. Kuwata, S. Grinstein, S. L. Schmid, and G. Danuser, Robust singleparticle tracking in live-cell time-lapse sequences, Nature Methods 5, 695 (2008).

