## Supplemental Information for "Active surface waves drive rippling in *Myxococcus xanthus* colonies"

Aaron R. Bourque*
*Lewis-Sigler Institute for Integrative Genomics, Princeton University, Princeton, NJ 08544, USA*

Peter A. E. Hampshire*
*Max Planck Institute for the Physics of Complex Systems, Dresden, Germany and*
*Center for Systems Biology Dresden, Dresden, Germany*

Ricard Alert†
*Max Planck Institute for the Physics of Complex Systems, Dresden, Germany*
*Center for Systems Biology Dresden, Dresden, Germany*
*Cluster of Excellence Physics of Life, TU Dresden, Dresden, Germany*
*Departament de Física de la Matèria Condensada, Universitat de Barcelona, Barcelona, Spain*
*Universitat de Barcelona Institute of Complex Systems (UBICS), Barcelona, Spain and*
*Institució Catalana de Recerca i Estudis Avançats (ICREA), Barcelona, Spain*

Joshua W. Shaevitz‡
*Joseph Henry Laboratories of Physics, Princeton University, Princeton, NJ 08544, USA and*
*Lewis-Sigler Institute for Integrative Genomics, Princeton University, Princeton, NJ 08544, USA*
(Dated: November 28, 2025)

* These authors contributed equally to this work.
†
‡

### Methods

#### I. COLONY-WIDE IMAGING OF *M. XANTHUS* RIPPLES

For brightfield imaging of whole rippling colonies of *M. xanthus*, we constructed a custom-built brightfield microscope consisting of a Basler Ace acA3088-57uc camera with a Computar MLH-10X manual zoom lens attached. A 60 mm petri dish containing a rippling colony was attached to a plate holder behind a NE40A-A absorptive ND filter and backlit by a QTH10 Thorlabs light source. The field of view of images is 3 mm x 5 mm, and images were acquired every 20 seconds for a period of 6 hours.

#### II. HEIGHT CHARACTERIZATION OF *M. XANTHUS* COLONIES

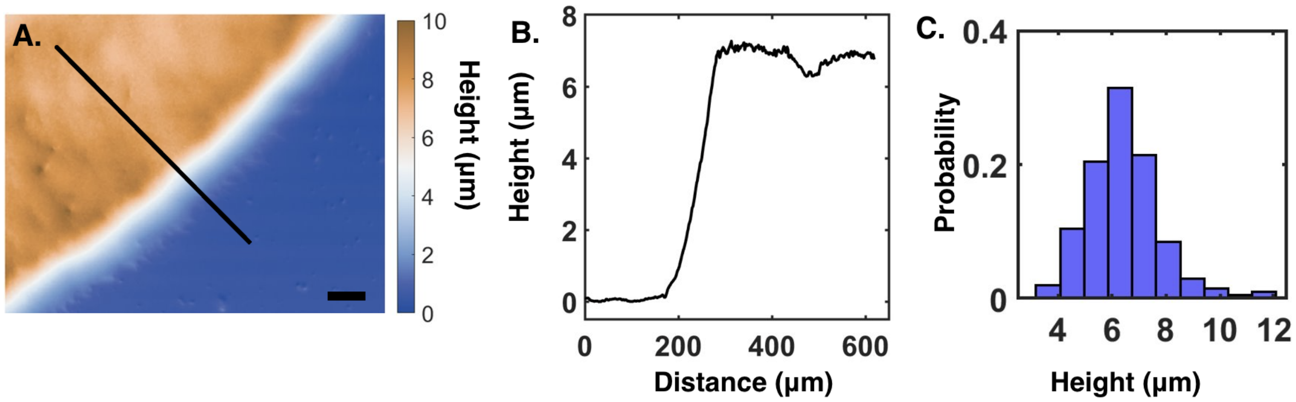


FIG. S1. Measurement of *M. xanthus* colony relative to substrate a) Height map of a colony *M. xanthus* on 1.5% agarose substrate prior to the formation of ripples. Scale bar = 100 μm. b) Height recorded along the line profile depicted in a). c) Distribution of average heights of *M. xanthus* colonies.

The Keyence VK-X1000 microscope is a laser scanning surface profilometer that illuminates rippling samples with a 661 nm laser and measures the reflectance of the laser at various sample heights to resolve a laser image. The height of the maximum intensity of the laser reflectance is recorded for each pixel for each image to resolve a height map of regions of regions of interest within a sample. The zero height for a time series is determined to be the pixel with the lowest height for all frames.

To estimate the absolute height of ripples and colony relative to the substrate, we conduct the rippling assay on 1.5% agarose substrates as described in Methods and Materials. The edge of the of the substrate is set to be the region of interest, which thus resolves a height map of the bare substrate, the colony boundary, and a portion of the rippling colony. The height is determined to be the zero height for the image, and line profiles are constructed from the substrate into the colony. Average height of a colony that is above the substrate is recorded. The measurement is repeated for a colony 10 times, where a line profile is radially distributed around the perimeter of the colony.

#### III. RHEOLOGICAL MEASUREMENTS OF HYDROGEL SUBSTRATES

The rheological properties of the agarose and phytagel substrates used in rippling experiments were calculated using an Anton Paar MCR 501 Rheometer [1]. CF medium containing the hydrogel gelling agents were melted to a temperature of 100 °C, and 1.5 mL of the molten medium was pipetted onto the rheometer stage heated at 80 °C. An Anton Paar Parallel-plate measuring probe (ISO 6721-10 and DIN 53019) was lowered to the molten media to a gap height of 1 mm. The stage heating was then deactivated, and the hydrogel was let cured at room temperature for 15 minutes.

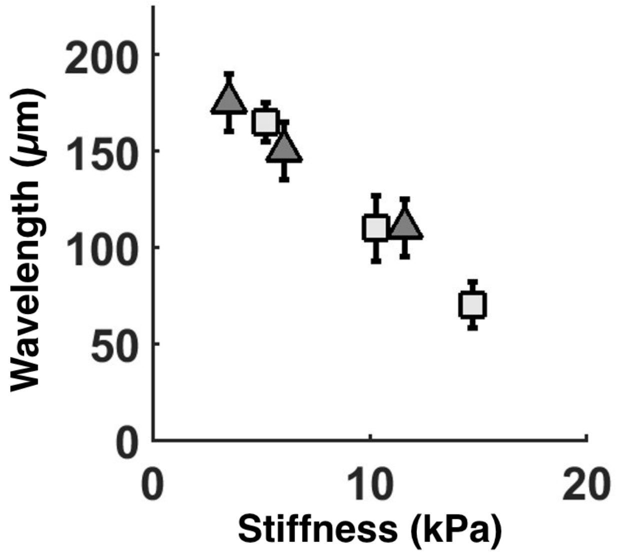


FIG. S2. Measurement of wavelength of rippling *M. xanthus* colonies on agarose (triangle) and phytagel (square) substrates.

An amplitude sweep test was then conducted at 22 °C at 0.1% strain for 8 cycles. A rheological amplitude sweep test measures the viscoelastic properties of a material by applying oscillatory shear with increasing strain amplitudes. This test helps identify the linear viscoelastic region (LVR) and determine the strain level where the material's structure starts to break down, indicating its transition from elastic to viscous behavior. The storage moduli reported for the different hydrogels was calculated by averaging the values measured for the various angular frequencies within the LVR. The reported storage moduli and corresponding rippling wavelengths are displayed in Fig. S3.

#### IV. TRACKING OF INDIVIDUAL CELLS VIA CONFOCAL IMAGING

| **Gelling Agent** | **Substrate Stiffness (kPa)** | **Speed ($\mu$m/s)** | **Standard Error ($\mu$m)** |
|---|---|---|---|
| Agarose | 1.1612 | 4.5656 | 0.1809 |
| Agarose | 0.6065 | 3.5035 | 0.2216 |
| Agarose | 0.3520 | 1.9589 | 0.2868 |
| Phytagel | 1.4738 | 3.5666 | 0.2241 |
| Phytagel | 1.0271 | 3.4639 | 0.1681 |
| Phytagel | 0.5191 | 3.9096 | 0.2318 |

In this study, two strains of *M. xanthus* were used in addition to the wild-type strain, DK1622. DK10547 has GFP transcriptionally fused to the *PilA* promoter, and it has been used in previous studies of rippling in *M. xanthus.* Furthermore, LS3908 expresses Td-Tomato and is grown with 1 mM IPTG in overnight cultures as described in Cotter et al [2]. Colonies containing entirely consisting of either DK10547 or LS3908 were observed to engage in rippling behavior. Furthermore, both colonies of these strains were measured to have the same rippling wavelength at 1.5% agarose. These strains are used for individual cell tracking in confocal imaging experiments, and motility values can be found in Table I.

#### V. CALCULATION OF THE RIPPLING WAVELENGTH VIA THE 2D FOURIER TRANSFORM

The 2D Fourier Transform was calculated on mean-centered height map data collected from the Keyence VK-X1000 microscope by calculating the fast Fourier Transform for the rows and columns of the image of size M×N. This results in the power spectrum of the image, an M×N-sized matrix in which each element represents spatial frequencies in $k$-space. The power spectrum is shifted such that the DC component of the signal is at the center of the matrix, resulting in two peaks that represent the dominant spatial frequency composing the waves (see Supplement). To calculate the rippling wavelength from this, we calculate the static structure factor by averaging the signal of the power spectrum of identical values of $k$. A log-normal distribution is then fitted for each time point of the duration of the rippling behavior. The mean of the fitted curve is determined to be the rippling wavelength for the image.

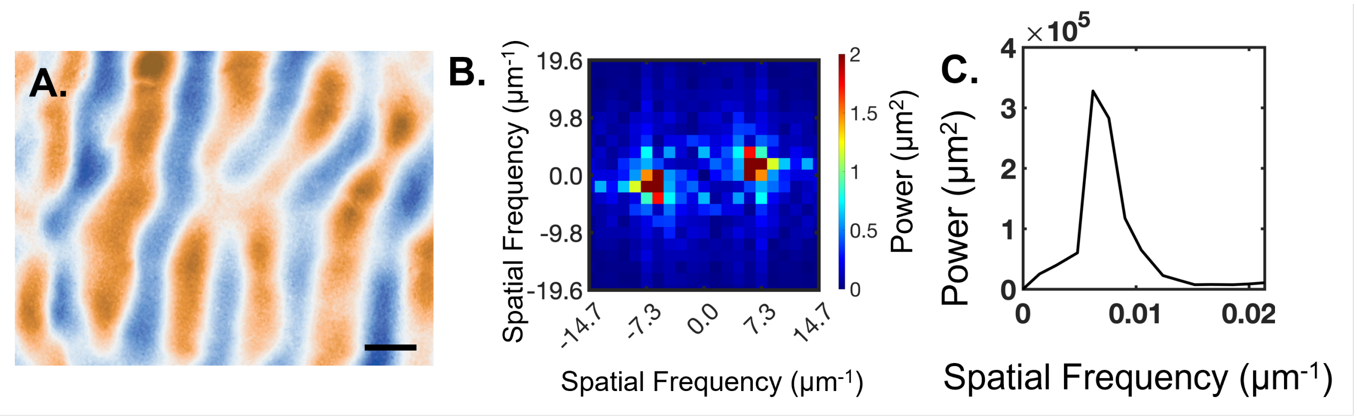


FIG. S3. 2D Fourier Transformation for calculation of the rippling wavelength. a) Height map of a rippling region of *M. xanthus* on 1.5% agarose substrate. Scale bar = 85 μm. b) Power spectrum of the image in a). The two peaks in the image depict the rippling wavelength. This is calculated by conducting the 2D Fourier transform of the image as described. The DC component of the signal is shifted to be in the center of the image. c). Averaged signal for each frequency from the image in b). The single peak is the rippling wavelength.

#### VI. CREST WIDTH MEASUREMENT

In models that describe accordion wave behavior of *M. xanthus*, the crest width of ripples is measured by recording the distance from one end of the density band to the other. In other words, the bare substrate marks the beginning and end of the width measurement. However, this approach is less straightforward when waves are three-dimensional in structure with cells in between them. To measure crest width the full width at half maximum for individual crests are measured across the height maps resolved from the Keyence VK-X1000 datasets, Fig. S4a, b.

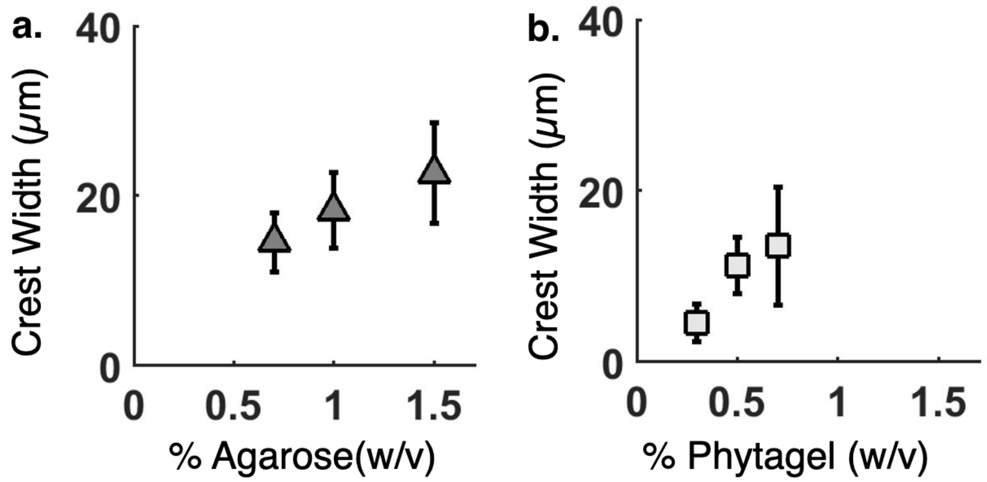


FIG. S4. Crest width measurements for rippling colonies of *M. xanthus*. a) Crest width measurement for agarose substrates. b) Crest width measurement for phytagel substrates.

#### VII. MEASUREMENTS OF MENISCUS WIDTH ON DIFFERENT SUBSTRATES

The meniscus width was measured by recording height profiles of individual cells on agarose of and phytagel substrates of varying stiffeness. First, DK16122 cells were grown overnight in CTTYE medium. 5 mL agarose and phytagel plates were prepared and left to dry uncovered for three hours. The plates were then submerged in 4 mL of CTTYE medium and incubated at 32 C for 4 hours. The *M. xanthus* cells were spun down and resuspended in CF medium to an optical density of 0.1. The CTTYE medium was then removed from the plates, and 10 µL of the prepared cells were incoulated onto the plates, and left to dry uncovered for an additional hour. The Keyence VK-X1000 microscope was then used to locate individual cells on each plate with the 100x magnification objective. Height maps of the individual cells were then measured to view the height profile across each cell. To calculate the meniscus length, the measured height profile was fitted to an exponential decay $h(x) = Ae^{-x/\ell_c} + c$, as described in Black et al. [3]. The fitted decay length, $\ell_c$, is determined to be the meniscus width.

#### VIII. MEASUREMENT OF SURFACE TENSION FOR LIQUID MEDIUM

Surface tension for CTTYE medium and water was measured via a custom-built pendant drop tensiometer. The tensiometer consists of a Basler Ace acA3088-57uc camera with a Computar MLH-10X manual zoom lens attached. Images of pendant drops were captured from a backlit, blunt-end needle (0.52 mm diameter) attached to a dispensing syringe. CTTYE medium containing TWN-80 surfactant was loaded into into the syringe, and pendant droplets were manually expelled from the syringe into the needle. Each trial is an individual pendant droplet that hangs from the needle. Images were processed using OpenDrop [4]. The surface tension of water is measured to be 73 ± 0.94 mN/m across 32 trials.

### Theory

#### IX. THEORY OF ACTIVE SURFACE WAVES

##### A. Active nematic hydrodynamics

As introduced in the Main Text, we consider a two-dimensional vertical cross-section of the bacterial colony (Fig. 3a), which we model as an incompressible, extensile active nematic fluid. The fluid is semi-infinite, with the colony-air interface at $y = h(x,t)$. Within the bulk, the fluid satisfies force balance, given by

$$0 = -\partial_\alpha P + \partial_\beta(\sigma_{\alpha\beta} + \sigma^{\mathrm{a}}_{\alpha\beta}). \tag{S1}$$

The pressure $P$ enforces the incompressibility condition $\boldsymbol{\nabla}\cdot\boldsymbol{v} = 0$, where $\boldsymbol{v}$ is the velocity field of the fluid. Respectively, $\sigma_{\alpha\beta}$ is the symmetric part of the deviatoric stress tensor, given by

$$\sigma_{\alpha\beta} = 2\eta u_{\alpha\beta} - \zeta Q_{\alpha\beta} + \frac{\nu}{2}(n_\alpha h_\beta + n_\beta h_\alpha - n_\gamma h_\gamma \delta_{\alpha\beta}), \tag{S2}$$

which includes (i) the viscous stress $2\eta u_{\alpha\beta}$, where $\eta$ is the viscosity and $u_{\alpha\beta} = (\partial_\alpha v_\beta + \partial_\beta v_\alpha)/2$ is the symmetric part of the strain rate tensor; (ii) the active nematic stress $-\zeta Q_{\alpha\beta}$, where $\zeta$ is the active stress coefficient and $Q_{\alpha\beta} = n_\alpha n_\beta - \delta_{\alpha\beta}/2$ is the nematic orientation tensor, where $\hat{\boldsymbol{n}} = (\cos\theta, \sin\theta)$ is the director field, $\theta$ is the director angle, and $\delta_{\alpha\beta}$ is the Kronecker delta; and (iii) flow-alignment stress $\frac{\nu}{2}(n_\alpha h_\beta + n_\beta h_\alpha - n_\gamma h_\gamma \delta_{\alpha\beta})$, where $\nu$ is the flow-alignment coefficient and $h_\alpha$ is the molecular field, which, in the one-constant approximation, is given by $h_\alpha = -\delta F/\delta n_\alpha = K\nabla^2 n_\alpha$, where $K$ is the Frank elastic constant and $F = K/2\int[(\boldsymbol{\nabla}\cdot\hat{\boldsymbol{n}})^2 + (\boldsymbol{\nabla}\times\hat{\boldsymbol{n}})^2]\,\mathrm{d}^2\boldsymbol{r}$ is the free energy of nematic distortions. We ignore the Ericksen stress, which is higher order in gradients of the director angle $\theta$. Finally, the last term in Eq. (S1) arises from the antisymmetric part of the deviatoric stress tensor, given by

$$\sigma^{\mathrm{a}}_{\alpha\beta} = \frac{1}{2}(n_\alpha h_\beta - n_\beta h_\alpha). \tag{S3}$$

The force balance condition implies that nematic distortions produce flows. In turn, the flows affect the nematic orientation, as captured in the following dynamics of the director:

$$\partial_t\hat{\boldsymbol{n}} + \boldsymbol{v}\cdot\boldsymbol{\nabla}\hat{\boldsymbol{n}} - \frac{1}{2}\boldsymbol{\omega}\times\hat{\boldsymbol{n}} = \frac{K}{\gamma_1}\nabla^2\hat{\boldsymbol{n}} - \nu\boldsymbol{u}\cdot\hat{\boldsymbol{n}}, \tag{S4}$$

which is Eq. 2 in the Main Text. Through this equation, the director varies via: (i) advection $\boldsymbol{v}\cdot\boldsymbol{\nabla}\hat{\boldsymbol{n}}$, (ii) corotation $-\frac{1}{2}\boldsymbol{\omega}\times\hat{\boldsymbol{n}}$, where $\boldsymbol{\omega}=\boldsymbol{\nabla}\times\boldsymbol{v}$ is the vorticity; (iii) flow alignment $-\nu\boldsymbol{u}\cdot\hat{\boldsymbol{n}}$; and (iv) elastic relaxation $\frac{K}{\gamma_1}\nabla^2\hat{\boldsymbol{n}}$, where $\gamma_1$ is the rotational viscosity.

##### B. Interface dynamics

Next we consider the dynamics at the colony-air interface. Firstly, the interface at $y=h(x,t)$ simply moves with the fluid. This means that we can write a dynamical equation for the height field by considering an element of fluid at the interface. This element has height $h_\mathrm{e}=h(x_\mathrm{e}(t),t)$, where $x_\mathrm{e}(t)$ is the $x$ coordinate of the element as a function of time. Taking the total derivative with respect to time gives

$$\frac{\mathrm{d}h_\mathrm{e}}{\mathrm{d}t}=\frac{\partial h(x_\mathrm{e}(t),t)}{\partial x_\mathrm{e}}\frac{\mathrm{d}x_\mathrm{e}}{\mathrm{d}t}+\frac{\partial h(x_\mathrm{e}(t),t)}{\partial t}. \tag{S5}$$

The velocity of the element is the same as the velocity of the fluid, and hence we have $\mathrm{d}h_\mathrm{e}/\mathrm{d}t=v_y(y=h)$ and $\mathrm{d}x_\mathrm{e}/\mathrm{d}t=v_x(y=h)$. Finally, by considering the element to have position $x_\mathrm{e}=x$, we can replace $x_\mathrm{e}$ with $x$ and rearrange to obtain

$$\partial_t h=v_y(y=h)-v_x(y=h)\partial_x h. \tag{S6}$$

Moreover, the flows at the interface are subject to two boundary conditions. First, we impose a no shear stress boundary condition:

$$\left[\sigma^{\mathrm{tot}}_{\alpha\beta}t_\alpha p_\beta\right]_{y=h}=0, \tag{S7}$$

where $\sigma^{\mathrm{tot}}_{\alpha\beta}=-P\delta_{\alpha\beta}+\sigma_{\alpha\beta}+\sigma^{\mathrm{a}}_{\alpha\beta}$ is the total stress, $\hat{\boldsymbol{t}}$ is the vector tangential to the surface, and $\hat{\boldsymbol{p}}$ is the normal to the surface pointing away from the bulk nematic (Fig. 3a). Additionally, we balance the normal component of the stress with a normal interfacial stress $\sigma^{\mathrm{int}}$:

$$\left[\sigma^{\mathrm{tot}}_{\alpha\beta}p_\alpha p_\beta\right]_{y=h}=\sigma^{\mathrm{int}}, \tag{S8}$$

which arises from the interfacial free energy, whose functional form, as derived in a previous study [3], is given by $\sigma^{\mathrm{int}}=-\delta F_{\mathrm{int}}/\delta h$, where $F_{\mathrm{int}}$ is the interfacial free energy. We take the interfacial free energy to be given by

$$F_{\mathrm{int}}=\int\mathrm{d}x\left[\gamma\sqrt{1+(\partial_x h)^2}+\frac{1}{2}bh^2\right], \tag{S9}$$

which includes contributions from the surface tension $\gamma$ and from a gravity-like restoring force, with coefficient $b$, which captures the energy penalty associated with extracting water from the underlying hydrogel substrate. We justify the contribution of the restoring force below. The interfacial energy Eq. (S9) defines the capillary length $\ell_\mathrm{c}=\sqrt{\gamma/b}$, below which the interface relaxation is dominated by surface tension, and above which it is dominated by the restoring force.

##### C. Heuristic justification of the interfacial restoring force

Here, we provide heuristic arguments to justify the interfacial restoring force $-bh$ that we use to capture the effects of the energy cost of extracting water from the substrate. Previous work showed that individual cells are surrounded by a meniscus of water, whose profile $h_\mathrm{m}(\boldsymbol{r})$ is determined by the Young-Laplace condition:

$$\gamma\, C[h_\mathrm{m}(\boldsymbol{r})]=P, \tag{S10}$$

where $\gamma$ is the surface tension of the water-air interface, $C=\nabla^2 h_\mathrm{m}/(1+|\boldsymbol{\nabla}h_\mathrm{m}|^2)^{3/2}$ is the interface curvature, and $P$ is the pressure in the meniscus [3]. This meniscus pressure is related to the osmotic pressure difference across the substrate of the hydrogel substrate, and thus it represents the energy per unit volume required to extract water from the substrate. Together with surface tension, through Eq. (S10), it defines the meniscus decay length $\ell_\mathrm{m}=\gamma/P$. Beyond menisci around individual cells, we now seek to obtain an equation for the coarse-grained height field $h(\boldsymbol{r})$ of the water-air interface in a cell colony. To this end, we first consider a colony with low cell density. In this case, the distance between cells is on average larger than the meniscus width, so that the menisci around each cell remain separate. Thus, if there is a total of $N$ cells in an area $A$, the average height of the water-air interface, $h_0 \equiv \langle h(\boldsymbol{r})\rangle_{\boldsymbol{r}}$, is given by

$$\langle h\rangle = \frac{NV}{A} = \rho_0 V, \tag{S11}$$

where $V = \int h_{\mathrm{m}}\,\mathrm{d}^2\boldsymbol{r}$ is the volume of water in a single-cell meniscus, and $\rho_0 = N/A$ is the average areal cell density. Thus, Eq. (S11) shows that the average water height increases linearly with the average cell density $\rho_0$ at low cell densities.

As the cell density increases, menisci start overlapping, and hence the dependence $\langle h\rangle(\rho_0)$ is no longer linear. For a close-packed cell layer, the water height tends to the cell height. However, as a second layer of cells starts forming, the (projected) areal cell density increases beyond close packing, and the water height increases again linearly with the cell density as menisci form around cells on the second layer. This trend continues for higher layers of cells. Therefore, for a multilayered system, the water height $h$ increases approximately linearly with the areal cell density $\rho$.

To capture this behavior, we propose the following force balance for the height field at mechanical equilibrium:

$$\gamma\nabla^2 h + F_{\mathrm{cell}}\rho - bh = 0. \tag{S12}$$

In addition to surface tension forces, the balance includes the force $F_{\mathrm{cell}}$ exerted by the cells on the water-air interface, and the restoring force $-bh$ that opposes changes in height around the average set by the cell density. The spatial average of Eq. (S12) gives $h_0 = F_{\mathrm{cell}}\rho_0/b$, thus recapitulating the linear height-density relationship explained above.

In the model presented in the Main Text, the forces exerted by the cells on the water-air interface are accounted for in the colony's flow field $\boldsymbol{v}$, which drives interface motion. The other two forces in Eq. (S12) are interfacial forces that we include in the boundary condition for the normal component of the stress at the interface, which in our cross-sectional geometry reads $[\hat{\boldsymbol{p}}\cdot\boldsymbol{\sigma}\cdot\hat{\boldsymbol{p}}]_{y=h} = \gamma\partial_x^2 h - bh$, as stated in the Main Text. Here, as in the Main Text, the unit vector $\hat{\boldsymbol{p}}$ indicates the direction perpendicular to the interface (Fig. 3a).

##### D. Linear stability analysis

Here, we analyse the linear stability of the flat-interface, homogeneous state defined by $\boldsymbol{v} = \boldsymbol{0}$, $\theta = 0$, and $h = 0$. Following previous studies [5, 6], we consider perturbations of all the variables by replacing each variable $X$ with $X_0 + \delta X$, where $X_0$ is the value of the variable in the flat-interface state and $\delta X$ is the perturbation, expanding up to first order in the perturbations, and obtaining the coupled linear dynamics of the perturbations of the interface height $\delta h(x,t)$ and the director angle at the interface $\delta\theta(x, y = 0, t)$.

Below, we work with the Fourier components of these fields, $\widetilde{\delta h}(k,t)$ and $\widetilde{\delta\theta}(k,y,t)$. For a function $f(x)$, the Fourier transform is defined as

$$\tilde{f}(k) := \int f(x)\mathrm{e}^{-ikx}\,\mathrm{d}x, \tag{S13}$$

where $k$ is the wave vector along the $x$ axis.

The linear equation for the dynamics of the height field is obtained by Taylor expanding the velocity components in Eq. (S6) about $y = 0$, substituting $h = \delta h$ and $\boldsymbol{v} = \delta\boldsymbol{v}$, and neglecting non-linear terms to obtain

$$\partial_t\delta h(x,t) = \delta v_y(x, y = 0, t). \tag{S14}$$

Taking the Fourier transform gives

$$\partial_t\widetilde{\delta h} = \widetilde{\delta v_y}(0), \tag{S15}$$

where we used the shorthand that $f(y) := f(k,y,t)$ for any function $f$.

To obtain the linear equation for the director angle we first need to rewrite Eq. (S4) in terms of the director angle. We first note that $\partial_t n_x = \partial_t\cos\theta = -\sin\theta\,\partial_t\theta$ and $\partial_t n_y = \partial_t\sin\theta = \cos\theta\,\partial_t\theta$, which means that $\partial_t\theta = -\sin\theta\,\partial_t n_x + \cos\theta\,\partial_t n_y$. Therefore, by using this expression and substituting $\partial_t n_x$ and $\partial_t n_y$ from Eq. (S4), we obtain

$$\partial_t\theta + \boldsymbol{v}\cdot\boldsymbol{\nabla}\theta - \frac{1}{2}(\partial_x v_y - \partial_y v_x) = \frac{K}{\gamma_1}\nabla^2\theta - \frac{\nu}{2}[\cos 2\theta\,(\partial_x v_y + \partial_y v_x) + \sin 2\theta\,(\partial_y v_y - \partial_x v_x)]. \tag{S16}$$

By substituting $\theta = \delta\theta$ and $\boldsymbol{v} = \delta\boldsymbol{v}$, Taylor expanding the functions of $\delta\theta$ about $\delta\theta = 0$, neglecting non-linear terms and taking the Fourier transform, we obtain the dynamics of the director angle in Fourier space as

$$\partial_t \widetilde{\delta\theta} = \frac{1}{2}(ik\widetilde{\delta v_y} - \partial_y \widetilde{\delta v_x}) - \frac{\nu}{2}(ik\widetilde{\delta v_y} + \partial_y \widetilde{\delta v_x}) - \frac{K}{\gamma_1}k^2\delta\theta + \frac{K}{\gamma_1}\partial_y^2\delta\theta. \tag{S17}$$

We can eliminate $\widetilde{\delta v_x}$ by substituting $\widetilde{\delta v_x} = i/k\partial_y\widetilde{\delta v_y}$, obtained by substituting $\boldsymbol{v} = \delta\boldsymbol{v}$ into the incompressibility condition, taking the Fourier transform and rearranging. Hence we obtain

$$\partial_t \widetilde{\delta\theta} = \frac{i}{2k}\left(k^2\widetilde{\delta v_y} - \partial_y^2\widetilde{\delta v_y}\right) - \nu\frac{i}{2k}\left(k^2\widetilde{\delta v_y} + \partial_y^2\widetilde{\delta v_y}\right) - \frac{K}{\gamma_1}k^2\widetilde{\delta\theta} + \frac{K}{\gamma_1}\partial_y^2\widetilde{\delta\theta}. \tag{S18}$$

Finally, we evaluate at $y = 0$. Here, following previous work [6], we assume that the director angle $\theta(x, y, t)$ varies much more slowly in the coordinate $y$ than the velocity component $v_y$ close to the colony-air interface. This assumption allows us to neglect $\partial_y^2\widetilde{\delta\theta}(0)$. This approximation ensures that the linear stability analysis is analytically tractable. Relaxing this assumption is an interesting direction that we leave for future work. Therefore, we obtain

$$\partial_t \widetilde{\delta\theta}(0) = \frac{i}{2k}\left(k^2\widetilde{\delta v_y}(0) - \partial_y^2\widetilde{\delta v_y}(0)\right) - \nu\frac{i}{2k}\left(k^2\widetilde{\delta v_y}(0) + \partial_y^2\widetilde{\delta v_y}(0)\right) - \frac{Kk^2}{\gamma_1}\widetilde{\delta\theta}(0). \tag{S19}$$

To obtain closed equations for the linearised dynamics Eqs. (S15) and (S19), we need to solve for the velocity component and its second derivative with respect to $y$ at the interface, $\widetilde{\delta v_y}(0)$ and $\partial_y^2\widetilde{\delta v_y}(0)$ respectively, in terms of the angle field at the interface $\widetilde{\delta\theta}(0)$ and the height field $\widetilde{\delta h}$.

##### 1. *Velocity solution*

To solve for the velocity component, we use the force balance equation Eq. (S1) with explicit expressions for the stresses in terms of the velocity and director angle. First, we substitute in $\theta = \delta\theta$, $\boldsymbol{v} = \delta\boldsymbol{v}$ and $P = P_0 + \delta P$, then Taylor expand the functions of $\delta\theta$ about $\delta\theta = 0$ and neglect non-linear terms to obtain

$$\boldsymbol{0} = -\boldsymbol{\nabla}\delta P + \eta\nabla^2\delta\boldsymbol{v} + \delta\boldsymbol{f}^{\text{nem}}, \tag{S20}$$

where we have introduced the linearised force density arising from the director distortion, $\delta f_\alpha^{\text{nem}} = \partial_\beta\delta\sigma_{\alpha\beta}^{\text{nem}}$, where $\delta\sigma_{\alpha\beta}^{\text{nem}}$ is the first-order contribution of the stress arising from director distortion given by

$$\delta\boldsymbol{\sigma}^{\text{nem}} = \begin{pmatrix} 0 & -\zeta\delta\theta + \frac{1}{2}K(\nu+1)\nabla^2\delta\theta \\ -\zeta\delta\theta + \frac{1}{2}K(\nu-1)\nabla^2\delta\theta & 0 \end{pmatrix}, \tag{S21}$$

Note, that we also used the fact that gradients in pressure of the flat-interface state are zero, i.e. $\boldsymbol{\nabla}P_0 = 0$, which can easily be derived by substituting $\theta = 0$, $\boldsymbol{v} = \boldsymbol{0}$ and $P = P_0$ into the force balance equation. Next, we take the Fourier transform along the $x$ direction of the linearised force balance equation, Eq. (S20), and the incompressibility condition. This leads to three equations for the velocity components and the pressure: $\widetilde{\delta v_x}$, $\widetilde{\delta v_y}$, and $\widetilde{\delta P}$. Eliminating $\widetilde{\delta v_x}$ and $\widetilde{\delta P}$, we obtain the following equation for $\widetilde{\delta v_y}$:

$$\left(\partial_y^2 - k^2\right)\left(\partial_y^2 - k^2\right)\widetilde{\delta v_y}(k, y, t) = \frac{ik}{\eta}\left(\partial_y\widetilde{\delta f_x^{\text{nem}}} - ik\widetilde{\delta f_y^{\text{nem}}}\right). \tag{S22}$$

where the Fourier transform of the linearised force density is given by $\widetilde{\delta f_\alpha^{\text{nem}}} = ik\widetilde{\delta\sigma_{\alpha x}^{\text{nem}}} + \partial_y\widetilde{\delta\sigma_{\alpha y}^{\text{nem}}}$, and the Fourier transform of the linearised stress is given by

$$\widetilde{\delta\boldsymbol{\sigma}^{\text{nem}}} = \begin{pmatrix} 0 & -\zeta\widetilde{\delta\theta} + \frac{1}{2}K(\nu+1)(-k^2 + \partial_y^2)\widetilde{\delta\theta} \\ -\zeta\widetilde{\delta\theta} + \frac{1}{2}K(\nu-1)(-k^2 + \partial_y^2)\widetilde{\delta\theta} & 0 \end{pmatrix}. \tag{S23}$$

Then, we linearise the boundary conditions, Eqs. (S7) and (S8). By Taylor-expanding the boundary conditions about $y = 0$, substituting the expressions of the tangential and normal vectors,

$$\hat{\boldsymbol{t}} = \frac{1}{\sqrt{1 + (\mathrm{d}h/\mathrm{d}x)^2}}\begin{pmatrix} 1 \\ \mathrm{d}h/\mathrm{d}x \end{pmatrix}, \qquad \hat{\boldsymbol{p}} = \frac{1}{\sqrt{1 + (\mathrm{d}h/\mathrm{d}x)^2}}\begin{pmatrix} -\mathrm{d}h/\mathrm{d}x \\ 1 \end{pmatrix} \tag{S24}$$

and $\sigma^{\rm tot}_{\alpha\beta} = \eta(\partial_\alpha v_\beta + \partial_\beta v_\alpha) - P\delta_{\alpha\beta} + \sigma^{\rm nem}_{\alpha\beta}$, where $\sigma^{\rm nem}_{\alpha\beta}$ is expressed explicitly in terms of the director angle, then substituting $\boldsymbol{v} = \delta\boldsymbol{v}$, $h = \delta h$, $\theta = \delta\theta$ and $P = P_0 + \delta P$, Taylor expanding the functions of $\delta\theta$ about $\delta\theta = 0$ and neglecting non-linear terms, we obtain

$$\left[\eta\left(\partial_x \delta v_y + \partial_y \delta v_x\right) + \delta\sigma^{\rm nem}_{xy} + \zeta\partial_x \delta h\right]_{y=0} = 0, \tag{S25}$$

and

$$\left[2\eta\partial_y\delta v_y - \delta P + \delta\sigma^{\rm nem}_{yy}\right]_{y=0} = \gamma\partial_x^2\delta h - b\delta h = \gamma\left(\partial_x^2\delta h - \ell_{\rm c}^{-2}\delta h\right). \tag{S26}$$

Note, that in Eq. (S26), we used the fact that $0 = -P_0 + \zeta/2$, which can easily be derived by substituting $P = P_0$, $\boldsymbol{v} = 0$, $\theta = 0$, $h = 0$, and $\hat{\boldsymbol{p}} = (0, 1)$ into Eq. (S8). As for the force balance equation, we take the Fourier transform and, in combination with the Fourier transform of the incompressibility condition and the $x$ component of the force balance equation, we eliminate $\widetilde{\delta P}$ and $\widetilde{\delta v_x}$ to obtain

$$\left[\eta\left(\partial_y^2 + k^2\right)\widetilde{\delta v_y} - ik\widetilde{\delta\sigma^{\rm nem}_{xy}} + \zeta k^2\widetilde{\delta h}\right]_{y=0} = 0 \tag{S27}$$

and

$$\left[\frac{\eta}{k^2}(\partial_y^3 - 3k^2\partial_y)\widetilde{\delta v_y} - \widetilde{\delta\sigma^{\rm nem}_{yy}} - \frac{i}{k}\widetilde{\delta f^{\rm nem}_x}\right]_{y=0} = \gamma(k^2 + \ell_{\rm c}^{-2})\widetilde{\delta h}. \tag{S28}$$

We write the solution to Eq. (S22) and the boundary conditions Eqs. (S27) and (S28) as the sum of the solution $\widetilde{\delta v_y}^{\rm h}$ to the homogeneous equation with the required boundary conditions and a particular solution $\widetilde{\delta v_y}^{\rm p}$ to the inhomogeneous equation with homogeneous boundary conditions:

$$\widetilde{\delta v_y} = \widetilde{\delta v_y}^{\rm h} + \widetilde{\delta v_y}^{\rm p}. \tag{S29}$$

The homogeneous equation has the solution

$$\widetilde{\delta v_y}^{\rm h} = Ae^{ky} + Bye^{ky}, \tag{S30}$$

where constants $A$ and $B$ are determined by the boundary conditions and given by

$$A = \frac{-\gamma k(1 + k^2\ell_{\rm c}^2)\widetilde{\delta h} - \ell_{\rm c}^2 k\widetilde{\delta\sigma^{\rm nem}_{yy}}(0) - i\ell_{\rm c}^2\widetilde{\delta f^{\rm nem}_x}(0)}{2\eta k^2\ell_{\rm c}^2}, \tag{S31a}$$

$$B = \frac{\gamma k(1 + k^2\ell_{\rm c}^2)\widetilde{\delta h} - \zeta\ell_{\rm c}^2 k^2\widetilde{\delta h} + i\ell_{\rm c}^2 k\widetilde{\delta\sigma^{\rm nem}_{xy}}(0) + \ell_{\rm c}^2 k\widetilde{\delta\sigma^{\rm nem}_{yy}}(0) + i\ell_{\rm c}^2\widetilde{\delta f^{\rm nem}_x}(0)}{2\eta k\ell_{\rm c}^2}. \tag{S31b}$$

Respectively, the inhomogeneous equation has the solution

$$\begin{aligned}\widetilde{\delta v_y}^{\rm p} = \frac{ik}{\eta}\Bigg\{&\int_{-\infty}^{y}{\rm d}y'\left[C_3e^{ky} + C_4e^{-ky} + C_5ye^{ky} + C_6ye^{-ky}\right]\left(\partial_{y'}\widetilde{\delta f^{\rm nem}_x}(y') - ik\widetilde{\delta f^{\rm nem}_y}(y')\right)\\ &+\int_{y}^{0}{\rm d}y'\left[C_1e^{ky} + C_2ye^{ky}\right]\left(\partial_{y'}\widetilde{\delta f^{\rm nem}_x}(y') - ik\widetilde{\delta f^{\rm nem}_y}(y')\right)\Bigg\},\end{aligned} \tag{S32}$$

where

$$C_1 = \frac{(1-ky')e^{ky'} + (1+ky')e^{-ky'}}{4k^3}, \tag{S33a}$$

$$C_2 = \frac{(-1+2ky')e^{ky'} - e^{-ky'}}{4k^2}, \tag{S33b}$$

$$C_3 = \frac{(1-ky')e^{ky'}}{4k^3}, \tag{S33c}$$

$$C_4 = \frac{(1-ky')e^{ky'}}{4k^3}, \tag{S33d}$$

$$C_5 = \frac{(-1+2ky')e^{ky'}}{4k^2}, \tag{S33e}$$

$$C_6 = \frac{e^{ky'}}{4k^2}. \tag{S33f}$$

To close the coupled dynamics of $\tilde{h}$ and $\tilde{\theta}(0)$ in Eqs. (S15) and (S19), we need to evaluate $\widetilde{\delta v_y}(0)$ and $\partial_y^2 \widetilde{\delta v_y}(0)$. Firstly, we evaluate Eqs. S29-S33 at $y=0$ to give

$$\widetilde{\delta v_y}(0) = \widetilde{\delta v_y}^{\mathrm{h}}(0) + \widetilde{\delta v_y}^{\mathrm{p}}(0), \tag{S34}$$

where the homogeneous part is

$$\widetilde{\delta v_y}^{\mathrm{h}}(0) = \frac{-\gamma k(1+k^2\ell_{\mathrm{c}}^2)\widetilde{\delta h} - \ell_{\mathrm{c}}^2 k \widetilde{\delta\sigma_{yy}^{\mathrm{nem}}}(0) - i\ell_{\mathrm{c}}^2 \widetilde{\delta f_x^{\mathrm{nem}}}(0)}{2\eta k^2 \ell_{\mathrm{c}}^2} \tag{S35}$$

and the inhomogeneous part is

$$\widetilde{\delta v_y}^{\mathrm{p}}(0) = \frac{i}{2\eta k^2}\left\{\int_{-\infty}^{0} \mathrm{d}y'\,(1-ky')\,e^{ky'}\left(\partial_{y'}\widetilde{\delta f_x^{\mathrm{nem}}}(y') - ik\widetilde{\delta f_y^{\mathrm{nem}}}(y')\right)\right\}. \tag{S36}$$

The $y$-component of the Fourier transform of the linearised force density is given by $\widetilde{\delta f_y^{\mathrm{nem}}} = ik\widetilde{\delta\sigma_{yx}^{\mathrm{nem}}} + \partial_y\widetilde{\delta\sigma_{yy}^{\mathrm{nem}}}$. Thus, the last factor in Eq. (S36) is $(\partial_y\widetilde{\delta f_x^{\mathrm{nem}}} - ik\widetilde{\delta f_y^{\mathrm{nem}}}) = \partial_y(\widetilde{\delta f_x^{\mathrm{nem}}} - ik\widetilde{\delta\sigma_{yy}^{\mathrm{nem}}}) + k^2\widetilde{\delta\sigma_{yx}^{\mathrm{nem}}}$, and using integration by parts we rewrite Eq. (S36) as

$$\widetilde{\delta v_y}^{\mathrm{p}}(0) = \frac{i}{2\eta k^2}\left\{\widetilde{\delta f_x^{\mathrm{nem}}}(0) - ik\widetilde{\delta\sigma_{yy}^{\mathrm{nem}}}(0) \right.$$
$$\left. + k^2\int_{-\infty}^{0}\mathrm{d}y' y' e^{ky'}\left(\widetilde{\delta f_x^{\mathrm{nem}}}(y') - ik\widetilde{\delta\sigma_{yy}^{\mathrm{nem}}}(y')\right) + k^2\int_{-\infty}^{0}\mathrm{d}y'\,(1-ky')\,e^{ky'}\widetilde{\delta\sigma_{yx}^{\mathrm{nem}}}(y')\right\}. \tag{S37}$$

Then, the $x$ component of the Fourier transform of the linearised force density is $\widetilde{\delta f_x^{\mathrm{nem}}} = ik\widetilde{\delta\sigma_{xx}^{\mathrm{nem}}} + \partial_y\widetilde{\delta\sigma_{xy}^{\mathrm{nem}}}$. Using it in the equation above and integrating by parts gives

$$\widetilde{\delta v_y}^{\mathrm{p}}(0) = \frac{i}{2\eta k^2}\left\{\widetilde{\delta f_x^{\mathrm{nem}}}(0) - ik\widetilde{\delta\sigma_{yy}^{\mathrm{nem}}}(0) + k^2\int_{-\infty}^{0}\mathrm{d}y' y' e^{ky'} ik\left(\widetilde{\delta\sigma_{xx}^{\mathrm{nem}}}(y') - \widetilde{\delta\sigma_{yy}^{\mathrm{nem}}}(y')\right)\right.$$
$$\left. - k^2\int_{-\infty}^{0}\mathrm{d}y'\,(1+ky')\,e^{ky'}\widetilde{\delta\sigma_{xy}^{\mathrm{nem}}}(y') + k^2\int_{-\infty}^{0}\mathrm{d}y'\,(1-ky')\,e^{ky'}\widetilde{\delta\sigma_{yx}^{\mathrm{nem}}}(y')\right\}. \tag{S38}$$

Hence, combining Eqs. (S34), (S35) and (S38) gives

$$\widetilde{\delta v_y}(0) = -\frac{\gamma(1+k^2\ell_{\mathrm{c}}^2)\widetilde{\delta h}}{2\eta k \ell_{\mathrm{c}}^2} + \frac{i}{2\eta}\left\{ik\int_{-\infty}^{0}\mathrm{d}y' y' e^{ky'}\left(\widetilde{\delta\sigma_{xx}^{\mathrm{nem}}}(y') - \widetilde{\delta\sigma_{yy}^{\mathrm{nem}}}(y')\right)\right.$$
$$\left. - \int_{-\infty}^{0}\mathrm{d}y'\,(1+ky')\,e^{ky'}\widetilde{\delta\sigma_{xy}^{\mathrm{nem}}}(y') + \int_{-\infty}^{0}\mathrm{d}y'\,(1-ky')\,e^{ky'}\widetilde{\delta\sigma_{yx}^{\mathrm{nem}}}(y')\right\}. \tag{S39}$$

We next introduce Eq. (S23) to obtain

$$\widetilde{\delta v_y}(0) = -\frac{\gamma(1+k^2\ell_{\rm c}^2)\widetilde{\delta h}}{2\eta k\ell_{\rm c}^2} \tag{S40}$$

$$+\frac{ik\zeta}{\eta}\int_{-\infty}^{0}{\rm d}y' y' e^{ky'}\widetilde{\delta\theta} + \frac{ik^3\nu K}{2\eta}\int_{-\infty}^{0}{\rm d}y' y' e^{ky'}\left(1-\frac{1}{k^2}\partial_{y'}^2\right)\widetilde{\delta\theta} + \frac{ik^2 K}{2\eta}\int_{-\infty}^{0}{\rm d}y' e^{ky'}\left(1-\frac{1}{k^2}\partial_{y'}^2\right)\widetilde{\delta\theta}. \tag{S41}$$

Here, we use again the approximation that the director angle varies slowly close to the interface to replace $\widetilde{\delta\theta}(y)$ in the integrand with $\widetilde{\delta\theta}(y=0)$. Expressed mathematically, this requires that the director angle is constant over a distance up to the order of $1/k$ from the interface, because the decay length of the Green's function for the velocity is $1/k$. With this approximation, we obtain

$$\widetilde{\delta v_y}(0) = -\frac{\gamma(1+k^2\ell_{\rm c}^2)\widetilde{\delta h}}{2\eta k\ell_{\rm c}^2} \tag{S42}$$

$$+\frac{ik\zeta\widetilde{\delta\theta}(0)}{\eta}\int_{-\infty}^{0}{\rm d}y' y' e^{ky'} + \frac{ik^3\nu K\widetilde{\delta\theta}(0)}{2\eta}\int_{-\infty}^{0}{\rm d}y' y' e^{ky'} + \frac{ik^2 K\widetilde{\delta\theta}(0)}{2\eta}\int_{-\infty}^{0}{\rm d}y' e^{ky'}. \tag{S43}$$

Finally, evaluating the integrals explicitly gives

$$\widetilde{\delta v_y}(0) = -\frac{\gamma(1+k^2\ell_{\rm c}^2)\widetilde{\delta h}}{2\eta k\ell_{\rm c}^2} - \frac{i\zeta\widetilde{\delta\theta}(0)}{\eta k} - \frac{ik\nu K\widetilde{\delta\theta}(0)}{2\eta} + \frac{ikK\widetilde{\delta\theta}(0)}{2\eta}, \tag{S44}$$

which already specifies the vertical component of the velocity at the interface in terms of the height and angle perturbations.

We now proceed similarly to calculate $\partial_y^2\widetilde{\delta v_y}(0)$ as

$$\partial_y^2\widetilde{\delta v_y}(0) = \partial_y^2\widetilde{\delta v_y}^{\rm h}(0) + \partial_y^2\widetilde{\delta v_y}^{\rm p}(0). \tag{S45}$$

Using the results above for the homogeneous and the inhomogeneous parts of the velocity solution, we obtain

$$\partial_y^2\widetilde{\delta v_y}^{\rm h}(0) = \frac{\gamma k(1+k^2\ell_{\rm c}^2)\widetilde{\delta h} - 2\zeta\ell_{\rm c}^2 k^2\widetilde{\delta h} + 2i\ell_{\rm c}^2 k\widetilde{\delta\sigma_{xy}^{\rm nem}}(0) + \ell_{\rm c}^2 k\widetilde{\delta\sigma_{yy}^{\rm nem}}(0) + i\ell_{\rm c}^2\widetilde{\delta f_x^{\rm nem}}(0)}{2\eta\ell_{\rm c}^2} \tag{S46}$$

and

$$\partial_y^2\widetilde{\delta v_y}^{\rm p}(0) = \frac{i}{2\eta}\int_{-\infty}^{0}{\rm d}y'\,(-1+ky')\,e^{ky'}\left(\partial_{y'}\widetilde{\delta f_x^{\rm nem}}(y') - ik\widetilde{\delta f_y^{\rm nem}}(y')\right). \tag{S47}$$

Comparing this Eq. (S47) with Eqs. (S36) and (S38) gives

$$\partial_y^2\widetilde{\delta v_y}^{\rm p}(0) = -k^2\widetilde{\delta v_y}^{\rm p}(0)$$

$$= -\frac{i}{2\eta}\left\{\widetilde{\delta f_x^{\rm nem}}(0) - ik\widetilde{\delta\sigma_{yy}^{\rm nem}}(0) + k^2\int_{-\infty}^{0}{\rm d}y' y' e^{ky'} ik\left(\widetilde{\delta\sigma_{xx}^{\rm nem}}(y') - \widetilde{\delta\sigma_{yy}^{\rm nem}}(y')\right)\right.$$

$$\left. -k^2\int_{-\infty}^{0}{\rm d}y'\,(1+ky')\,e^{ky'}\widetilde{\delta\sigma_{xy}^{\rm nem}}(y') + k^2\int_{-\infty}^{0}{\rm d}y'\,(1-ky')\,e^{ky'}\widetilde{\delta\sigma_{yx}^{\rm nem}}(y')\right\}. \tag{S48}$$

Hence, combining Eqs. (S46) and (S48) gives

$$\partial_y^2\widetilde{\delta v_y}(0) = \frac{\gamma k(1+k^2\ell_{\rm c}^2)\widetilde{\delta h} - 2\zeta\ell_{\rm c}^2 k^2\widetilde{\delta h} + 2i\ell_{\rm c}^2 k\widetilde{\delta\sigma_{xy}^{\rm nem}}(0)}{2\eta\ell_{\rm c}^2}$$

$$-\frac{ik^2}{2\eta}\left\{ik\int_{-\infty}^{0}{\rm d}y' y' e^{ky'}\left(\widetilde{\delta\sigma_{xx}^{\rm nem}}(y') - \widetilde{\delta\sigma_{yy}^{\rm nem}}(y')\right)\right.$$

$$\left. -\int_{-\infty}^{0}{\rm d}y'\,(1+ky')\,e^{ky'}\widetilde{\delta\sigma_{xy}^{\rm nem}}(y') + \int_{-\infty}^{0}{\rm d}y'\,(1-ky')\,e^{ky'}\widetilde{\delta\sigma_{yx}^{\rm nem}}(y')\right\}. \tag{S49}$$

Using Eq. (S23) and the assumption that the director angle varies slowly close to the interface as before, we arrive at

$$\partial_y^2\widetilde{\delta v_y}(0) = \frac{\gamma k(1+k^2\ell_{\rm c}^2)\widetilde{\delta h} - 2\zeta\ell_{\rm c}^2 k^2\widetilde{\delta h} - 2i\ell_{\rm c}^2\zeta k\widetilde{\delta\theta}(0) - i\ell_{\rm c}^2 K(\nu+1)k^3\widetilde{\delta\theta}(0)}{2\eta\ell_{\rm c}^2}$$
$$-\frac{i\zeta k^3\widetilde{\delta\theta}(0)}{\eta}\int_{-\infty}^{0}\mathrm{d}y' y' e^{ky'} - \frac{ik^5\nu K\widetilde{\delta\theta}(0)}{2\eta}\int_{-\infty}^{0}\mathrm{d}y' y' e^{ky'} - \frac{ik^4 K\widetilde{\delta\theta}(0)}{2\eta}\int_{-\infty}^{0}\mathrm{d}y' e^{ky'}. \tag{S50}$$

Finally, evaluating the integrals explicitly gives

$$\partial_y^2\widetilde{\delta v_y}(0) = \frac{[\gamma k(1+k^2\ell_{\rm c}^2) - 2\zeta\ell_{\rm c}^2 k^2]\widetilde{\delta h}}{2\eta\ell_{\rm c}^2} - \frac{ik^3 K\widetilde{\delta\theta}(0)}{\eta}, \tag{S51}$$

which gives $\partial_y^2\widetilde{\delta v_y}(0)$ in terms of the height and angle perturbations.

##### *2. Closed linear dynamics*

Substituting Eqs. (S44) and (S51) into Eqs. (S15) and (S19) gives the closed equations for the coupled linear dynamics of height and angle perturbations, which can be written as

$$\partial_t \begin{pmatrix} \widetilde{\delta h} \\ \widetilde{\delta\theta}(0) \end{pmatrix} = \boldsymbol{J}\begin{pmatrix} \widetilde{\delta h} \\ \widetilde{\delta\theta}(0) \end{pmatrix} \tag{S52}$$

where the Jacobian matrix is given by

$$\boldsymbol{J} = \begin{pmatrix} -\dfrac{\gamma(1+k^2\ell_{\rm c}^2)}{2\eta k\ell_{\rm c}^2} & -\dfrac{i\zeta}{\eta k} + \dfrac{ikK(1-\nu)}{2\eta} \\ -\dfrac{i\gamma(1+k^2\ell_{\rm c}^2)}{2\eta\ell_{\rm c}^2} + \dfrac{i\zeta(1+\nu)k}{2\eta} & \dfrac{\zeta(1-\nu)}{2\eta} - \dfrac{k^2K(3+\nu^2)}{4\eta} - \dfrac{k^2K}{\gamma_1} \end{pmatrix} \tag{S53}$$

#### E. Selected wavelength and frequency

As explained in the Main Text, these dynamics can give rise to waves. In particular, the theory predicts that, above a critical activity, there is a range of unstable, oscillatory modes (shown with positive growth rates and corresponding oscillation frequencies in Fig. 3b). The growth rates of these modes is given by $g(k) = \mathrm{Tr}\,\boldsymbol{J}/2$ and the oscillation frequency is $\omega(k) = \sqrt{-\mathrm{Tr}(\boldsymbol{J})^2/4 + \det(\boldsymbol{J})}$. To compare to the experimental observations, we predict that waves occur with the selected wavelength corresponding to the wave vector $k$ with the highest growth rate $g$. For simplicity, here we assume that the Frank elastic constant is small enough to neglect its contribution to the growth rate. In this limit, the growth rate is given by

$$g(k) = \frac{1}{4\eta}\left[\zeta(1-\nu) - \frac{\gamma(1+k^2\ell_{\rm c}^2)}{k\ell_{\rm c}^2}\right]. \tag{S54}$$

The peak growth rate occurs when $\mathrm{d}g/\mathrm{d}k = 0$, which yields a selected wave vector of

$$k_{\rm s} = 1/\ell_{\rm c}, \tag{S55}$$

and therefore the selected wavelength is

$$\lambda_{\rm s} = 2\pi\ell_{\rm c} = 2\pi\sqrt{\frac{\gamma}{b}}. \tag{S56}$$

By substituting Eq. (S55) into the expression for the oscillation frequency, $\omega(k) = \sqrt{-\mathrm{Tr}(\boldsymbol{J})^2/4 + \det(\boldsymbol{J})}$, we obtain the selected frequency as

| Parameter | Description | Estimate |
|---|---|---|
| $\ell_c$ | Capillary length | 16 μm |
| $\gamma$ | Surface tension | 65 mN/m |
| $\nu$ | Flow alignment coefficient | -1.1 |
| $\zeta$ | Active stress coefficient | 4 kPa |
| $K$ | Frank elastic constant | 2 kPa $\mu$m$^2$ |
| $\eta$ | Viscosity | 300 Pa min |
| $\gamma_1$ | Rotational viscosity | 300 Pa min |

TABLE I. Model parameter estimates.

$$\omega_{\rm s} = \frac{1}{8}\sqrt{\frac{-4\ell_{\rm c}^2(\ell_{\rm c}(\nu+3)\zeta-2\gamma)^2+4K\ell_{\rm c}\left(2\gamma\left(\nu^2+4\nu+3\right)-\ell_{\rm c}\left(\nu^3+3\nu^2+7\nu-11\right)\zeta\right)-K^2\left(\nu^2+7\right)^2}{\eta^2\ell_{\rm c}^4}}, \tag{S57}$$

which can equivalently be written in terms of the active time $\tau_{\rm a} = \eta/\zeta$ as

$$\omega_{\rm s} = \tau_{\rm a}^{-1} f({\rm Ca_A}, \ell_{\rm a}/\ell_{\rm c}, \nu). \tag{S58}$$

Here, the function $f$ has only dimensionless arguments: the active capiliary number ${\rm Ca_A} \equiv \zeta\ell_{\rm c}/\gamma$, the ratio of the active length $\ell_{\rm a} = \sqrt{K/\zeta}$ to the capillary length $\ell_{\rm c}$ and the flow-alignment coefficient $\nu$, and is given by

$$f({\rm Ca_A}, \ell_{\rm a}/\ell_{\rm c}, \nu) = \frac{1}{8}\sqrt{-4(\nu+3-2{\rm Ca_A^{-1}})^2+4\frac{\ell_{\rm a}^2}{\ell_{\rm c}^2}[2{\rm Ca_A^{-1}}(\nu^2+4\nu+3)-(\nu^3+3\nu^2+7\nu-11)]-\frac{\ell_{\rm a}^4}{\ell_{\rm c}^4}(\nu^2+7)^2}. \tag{S59}$$

##### F. Parameter estimates

Here, we estimate model parameter values based on the experimental results. We use the parameter estimates obtained here for the plot of the dispersion relation (Fig. 3b) in the Main Text. Table I shows the parameter estimates used for all plots unless stated explicitly otherwise. They are determined as follows.

Firstly, we use our measurements of the wavelength of the ripples, $\lambda_{\rm s} \sim 100$ μm and Eq. (S56), to estimate the capilliary length as $\ell_{\rm c} \sim 16$ μm. Additionally, we use our measurements of the surface tension of the colony $\gamma \approx 65$ kPa μm.

To make progress, we make three simplifying assumptions regarding the other parameters. First, we assume that the rotational viscosity is equal to the viscosity, i.e. $\gamma_1 = \eta$. Next, we take the flow-alignment coefficient to be $\nu \approx -1.1$, which is a typical value for liquid crystals made of flow-aligning rods ($\nu < -1$) [7]. Later, we assess the effect of the flow alignment on the dispersion relation (Fig. S6). Finally, we assume that the active stress coefficient is just above the value required for the growth rate to be positive at the selected wavelength. By setting $k = 1/\ell_{\rm c}$ in Eq. (S54), the condition $g(1/\ell_{\rm s}) > 0$ implies $\zeta > 2\gamma/[\ell_{\rm c}(1-\nu)]$, from which we estimate $\zeta \sim 4$ kPa. Later, we further quantify the effect of the active stress coefficient on the dispersion relation (Fig. S6).

In order to estimate the value of the Frank elastic constant, we determine how the dispersion relation changes as its value is increased, as shown in Fig. S5. We require that the growth rate is positive and there are oscillations at the selected wave vector $k = 1/\ell_{\rm c}$. At low Frank elastic constants, of $K \lesssim 1$ kPa μm$^2$, there are no oscillations at the selected wavelength, whereas at high Frank elastic constants, of $K \gtrsim 20$ kPa μm$^2$, the growth rate at the selected wavelength is negative. However, at the intermediate value $K = 2$ kPa μm$^2$, both conditions are satisfied, so we estimate the Frank elastic constant as $K \sim 2$ kPa μm$^2$.

Next, we use our measurements of the period of the ripples $T \approx 20$ min and the analytical expression for the oscillation frequency at the selected wave vector $\omega(k_{\rm s}) = \sqrt{-{\rm Tr}(\boldsymbol{J})^2/4 + \det(\boldsymbol{J})}|_{k=k_{\rm s}}$ to estimate the viscosity of the colony. This gives $\eta \sim 300$ Pa min.

Finally, we explore the roles of the flow-alignment coefficient $\nu$ and active stress coefficient $\zeta$ to further justify the estimates discussed above. We plot how the dispersion relation varies with these parameters in Fig. S6. When the flow-alignment coefficient has a small negative value ($\nu = -0.8$), the growth rate at the selected wavelength is negative. When the flow-alignment coefficient has a large negative value ($\nu = -1.4$), there are no oscillations at the selected wave vector. In contrast, for intermediate values such as $\nu = -1.1$, there are unstable oscillating modes consistent with the emergence of waves. Similarly, when the active stress coefficient is too low or too high, e.g. $\zeta = 3.5$ or 5 kPa respectively, there are no oscillations at the selected wave vector. This justifies our estimate of an intermediate value of $\zeta = 4$ kPa.

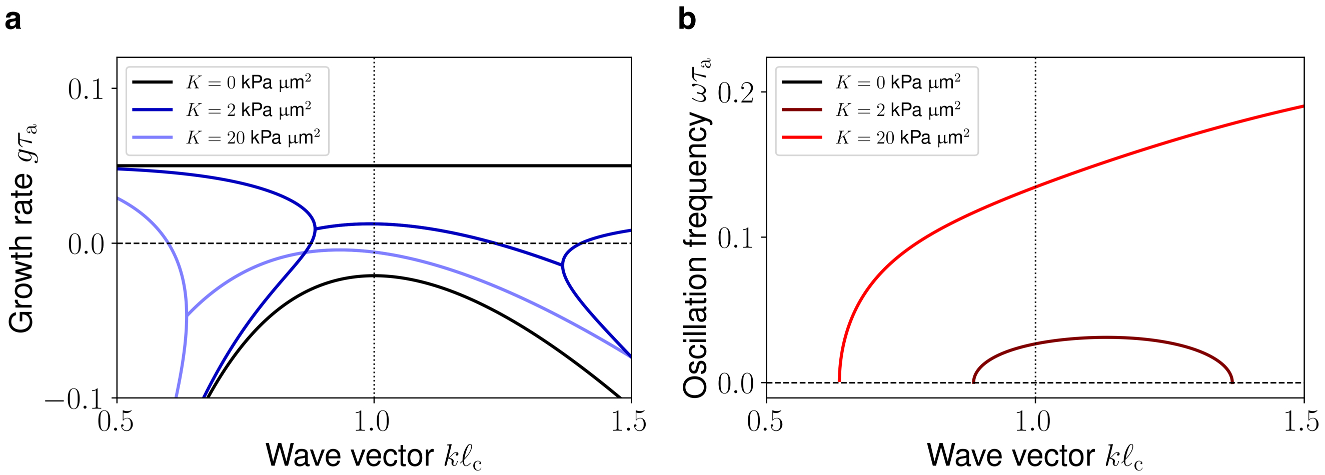


FIG. S5. **Role of Frank elasticity on the dispersion relation.** The growth rate $g$ (a) and oscillation frequency $\omega$ (b) of interfacial perturbations of wave vector $k$ for different values of the Frank elastic constant $K$. The regions with two branches of the growth rate feature modes with two real eigenvalues, whereas regions with a single branch of the growth rate feature modes with complex conjugate eigenvalues, which indicate the presence of oscillations with frequency given by the imaginary part. The vertical dotted line at $k = 1/\ell_c$ indicates the selected wave vector (see text).

[1] J. K. Nunes, H. Constantin, and H. A. Stone, Microfluidic tailoring of the two-dimensional morphology of crimped microfibers, Soft Matter **9**, 4227 (2013), publisher: Royal Society of Chemistry.

[2] C. R. Cotter, H.-B. Schüttler, O. A. Igoshin, and L. J. Shimkets, Data-driven modeling reveals cell behaviors controlling self-organization during myxococcus xanthus development, Proceedings of the National Academy of Sciences **114**, E4592 (2017).

[3] M. E. Black, C. Fei, R. Alert, N. S. Wingreen, and J. W. Shaevitz, Capillary interactions drive the self-organization of bacterial colonies, Nature Physics **21**, 1444 (2025).

[4] J. D. Berry, M. J. Neeson, R. R. Dagastine, D. Y. C. Chan, and R. F. Tabor, Measurement of surface and interfacial tension using pendant drop tensiometry, Journal of Colloid and Interface Science **454**, 226 (2015).

[5] R. Adkins, I. Kolvin, Z. You, S. Witthaus, M. C. Marchetti, and Z. Dogic, Dynamics of active liquid interfaces, Science **377**, 768 (2022).

[6] P. Gulati, F. Caballero, I. Kolvin, Z. You, and M. C. Marchetti, Traveling waves at the surface of active liquid crystals, Soft Matter **20**, 7703 (2024).

[7] S. A. Edwards and J. M. Yeomans, Spontaneous flow states in active nematics: A unified picture, Europhysics Letters **85**, 18008 (2009).

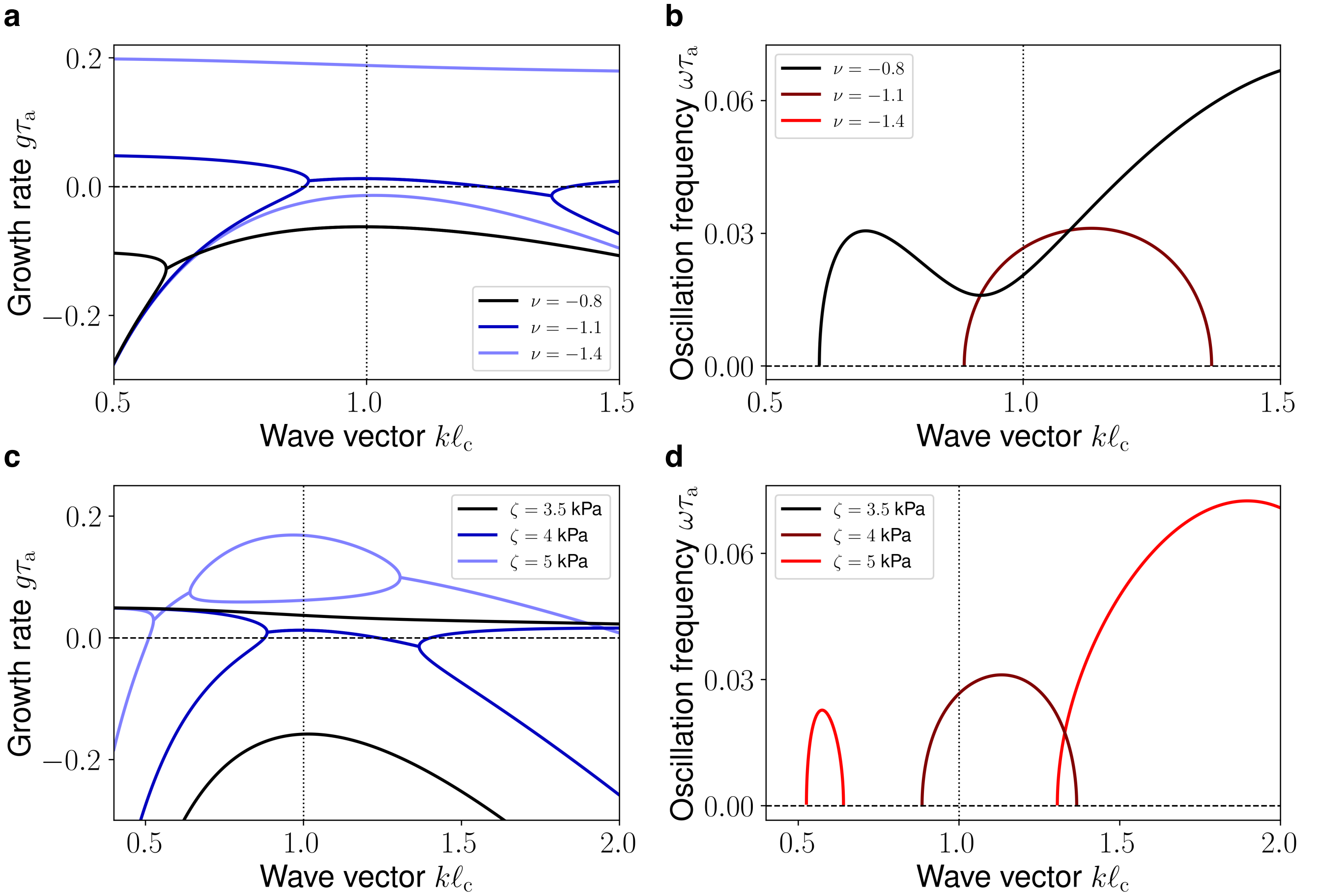


FIG. S6. **Role of flow alignment and activity on the dispersion relation.** The growth rate $g$ (a,c) and oscillation frequency $\omega$ (b,d) of interfacial perturbations of the wave vector $k$ for different values of the flow-alignment parameter $\nu$ (a,b) and of the active stress coefficient $\zeta$ (c,d). The regions with two branches of the growth rate feature modes with two real eigenvalues, whereas regions with a single branch of the growth rate feature modes with complex conjugate eigenvalues, which indicate the presence of oscillations with frequency given by the imaginary part. The vertical dotted line at $k = 1/\ell_c$ indicates the selected wave vector (see text).
